# The Calcitron: A Simple Neuron Model That Implements Many Learning Rules via the Calcium Control Hypothesis

**DOI:** 10.1101/2024.01.16.575890

**Authors:** Toviah Moldwin, Li Shay Azran, Idan Segev

**Affiliations:** Edmond and Lily Safra Center for Brain Sciences, The Hebrew University of Jerusalem; Department of Brain Sciences, Weizmann Institute of Science, Rehovot, Israel; Department of Neurobiology, The Hebrew University of Jerusalem

**Author notes:** **Correspondence** Toviah Moldwin.

**Keywords:** Synaptic weights, synaptic plasticity, calcium-based plasticity, neural plasticity, perceptron, machine learning, neural computation, Hebbian, homeostatic

## Abstract

Theoretical neuroscientists and machine learning researchers have proposed a variety of learning rules for linear neuron models to enable artificial neural networks to accomplish supervised and unsupervised learning tasks. It has not been clear, however, how these theoretically-derived rules relate to biological mechanisms of plasticity that exist in the brain, or how the brain might mechanistically implement different learning rules in different contexts and brain regions. Here, we show that the calcium control hypothesis, which relates plastic synaptic changes in the brain to calcium concentration [Ca^2+^] in dendritic spines, can reproduce a wide variety of learning rules, including some novel rules. We propose a simple, perceptron-like neuron model that has four sources of [Ca^2+^]: local (following the activation of an excitatory synapse and confined to that synapse), heterosynaptic (due to activity of adjacent synapses), postsynaptic spike-dependent, and supervisor-dependent. By specifying the plasticity thresholds and amount of calcium derived from each source, it is possible to implement Hebbian and anti-Hebbian rules, one-shot learning, perceptron learning, as well as a variety of novel learning rules.

## Introduction

Artificial neural networks (ANNs) have demonstrated a remarkable ability to solve both supervised and unsupervised learning tasks (Lecun et al., 2015). Because ANNs are inspired by a simple model of biological neurons and their synapses (McCulloch & Pitts, 1943), theoretical neuroscientists have used ANNs to explore questions about the brain. (Amit, 1992; Kriegeskorte, 2015; Memmesheimer et al., 2014; D. Rumelhart et al., 1986; Saxe et al., 2019). ANNs learn to solve a wide variety of problems by making use of “learning rules” whereby connection strengths between network nodes are modified (Hebb, 1949; Kohonen, 1982; Oja, 1982; Rosenblatt, 1958; D. E. Rumelhart et al., 1986). These learning rules are analogous to, and often inspired by, rules governing synaptic plasticity in biological neurons. However, it is often not apparent how biological plasticity mechanisms can result in the types of learning rules used in ANNs.

One of the dominant theories for how long-term synaptic plasticity operates in the brain is the calcium control hypothesis (Graupner & Brunel, 2012; J. Lisman, 1989; J. E. Lisman, 2001; J. Lisman & Goldring, 1988; Shouval et al., 2002, 2010). The calcium control hypothesis states that the magnitude and direction of change of synaptic strength is mediated by the intracellular calcium concentration ([Ca^2+^]) at the synapse. In the classic version of the calcium control hypothesis, at low levels of [Ca^2+^], the synapse is unaffected, at medium levels of [Ca^2+^], the synapse is depressed, and at high levels of [Ca^2+^], the synapse is potentiated. There is also some evidence that at very high levels of [Ca^2+^], all plastic mechanisms are turned off (Tigaret et al., 2016). This description of the relationship between calcium concentration and plasticity fits with experimental evidence from hippocampus and cortex (Artola et al., 1990; J. Lisman, 1989). In cerebellar Purkinje cells however, the calcium thresholds for potentiation and depression seem to be reversed; medium concentrations of [Ca^2+^] cause potentiation and high concentrations lead to depression (Coesmans et al., 2004; Piochon et al., 2016). While many downstream molecular mechanisms (most notably CaMKII and calcineurin) are involved in mediating plasticity (Citri & Malenka, 2008; Sanes & Lichtman, 1999), the calcium control hypothesis remains one of the most parsimonious and effective theories for how synaptic plasticity works in the brain. It remains to be explained, however, how calcium control of plasticity can result in the incredible learning abilities of the brain, especially if we work with the assumption that the brain operates in a manner similar to artificial neural networks.

In this work, we aim to bridge the gap between learning rules in artificial neurons and networks and the biological mechanisms of synaptic plasticity. We propose a simple, perceptron-like threshold-linear neuron model, the *calcitron*, that has four potential sources of local (synapse-specific) and global (common to all synapses) calcium. By adjusting the amount of calcium obtained from each calcium source and the calcium thresholds for plasticity, we show that it is possible to implement a wide variety of learning rules, such as Hebbian and anti-Hebbian learning, frequency-dependent plasticity, homeostatic plasticity, the perceptron learning rule, one-shot “writable” neural memory, and more. We thereby demonstrate that calcium control of synaptic plasticity can be a highly versatile mechanism which enables neurons to implement many different “programs” for modifying their synapses and storing information.

## Results

### The Calcitron Model

The calcitron is a simple neuron model, akin to a McCulloch and Pitts (M&P) neuron, or perceptron, which applies a transfer function to the weighted sum of its inputs. Formally we have:

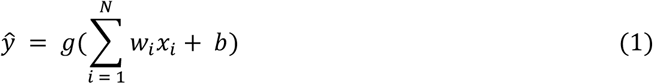

where *ŷ*is the output of the neuron, *g* is the transfer function, *w*_*i*_ is the weight of synapse *i, x*_*i*_ is the input to synapse *i, N* is the total number of synapses and *b* is a bias term. Depending on the particular use case, the transfer function *g* can be a simple threshold nonlinearity (e.g. a sign function), a sigmoid, a linear function, or any of the standard transfer functions used for artificial neural networks, so the set of possible outputs *ŷ*depend are contingent on the choice of transfer function. The weights *w*_*i*_ and inputs *x*_*i*_ of the calcitron are generally restricted to be non-negative to maintain fidelity to the experimental literature of calcium control hypothesis which mostly focuses on excitatory synapses (Graupner & Brunel, 2012; J. Lisman, 1989; J. E. Lisman, 2001; J. Lisman & Goldring, 1988; Shouval et al., 2002, 2010). In some contexts, the inputs *x*_*i*_ will be restricted to be binary (0 or 1); in other contexts, the inputs can be any positive real value (i.e., a rate model). The synaptic weights are also bounded between a minimum and maximum strength (*w*_*min*_ and *w*_*max*_, respectively) due to the nature of the calcium-based plasticity rule; this will be discussed further below. Because the weights and inputs are restricted in our model to be excitatory, the bias term *b* will generally be negative or 0, thus representing the aggregate inhibitory input to the neuron.

The calcitron has four sources of calcium. The first source of calcium is the local calcium due to the presynaptic input at each synapse, *C*_*local*_. The local calcium at synapse *i* is defined as:

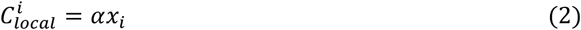

where *α* is a non-negative coefficient that determines the marginal increase in spine calcium for a unitary increase in input magnitude. Biologically, *C*_*local*_ can be thought of as the calcium that enters a dendritic spine receiving excitatory input through its NMDA receptors during synaptic stimulation. We note that according to the above formulation, the local calcium does *not* depend on the synaptic weight, *w*_*i*_, only on the synaptic input, *x*_*i*_. This decision is motivated by both biological and computational considerations. Biologically, synaptic strength is a consequence of AMPA receptor conductance (Citri & Malenka, 2008) while plasticity-inducing calcium primarily via NMDA receptors. Thus, while changing the synaptic weight (i.e., the number of AMPA receptors) will influence the somatic depolarization observed for a given presynaptic input, it will not necessarily change the calcium influx via the NMDA receptors. (It is true that the NMDA receptor’s conductance is also voltage-dependent (Jahr et al., 1990; Jahr & Stevens, 1990), so the increase of AMPA conductance of the synapse, as well as the weighted input from other synapses, can indirectly affect the calcium influx by depolarizing the neuron. We model the aggregate calcium influx due to input-dependent depolarization as heterosynaptic calcium, see next paragraph and **Discussion**). The weight-independence of the local calcium influx has the computational advantage of avoiding feedback loops – if calcium influx was weight-dependent, potentiating or depressing a synapse would change the synapse’s sensitivity to plasticity protocols.

The second source of intracellular calcium in the Calcitron is calcium that globally enters dendritic spines due to the aggregate activity of all nearby synaptic inputs, resulting in the activation of voltage-gated calcium channels (VGCCs) in regions of the dendrite that are sufficiently depolarized (J. E. Lisman, 2001; Moldwin, Kalmenson, et al., 2023) (and by increasing the conductance of NMDA at active synapses via NMDA’s voltage dependence, see **Discussion**). We call this calcium source heterosynaptic calcium, or *C*_*het*_. This calcium is responsible for heterosynaptic plasticity, i.e. plasticity that can be induced at non-activated synapses by presynaptic stimulation at other nearby synapses. While heterosynaptic plasticity is spatially sensitive, for simplicity we assume that synaptic activity is distributed uniformly on the neuron and we can thus approximate this calcium, *C*_*het*_, by looking at the aggregate activity from all synaptic inputs. We thus have

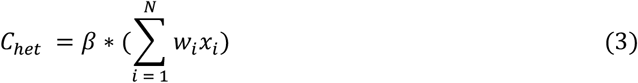

where *β* is a coefficient that determines how much calcium enters each spine due to the overall depolarization of the dendritic membrane and *x*_*i*_ and *w*_*i*_ are as in Eq. (1).

The third source of calcium, *C*_*BAP*_, comes from the backpropagating action potential, or BAP. When a neuron fires an action potential, the axonal/somatic spike backpropagates to the dendrites and depolarizes the dendritic membrane (Stuart & Sakmann, 1994), which can globally activate voltage-gated calcium channels at all spines. We model this as:

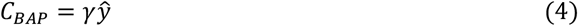

Where *γ* is a coefficient that determines the amount of calcium that enters the postsynaptic spines due to each spike. (Here we ignore timing effects of the postsynaptic spike relative to the timing of the input; for the purposes of the calcitron we assume that synaptic input and the spike it generates happen within a single time step. This assumption prevents the calcitron from implementing spike-timing dependent plasticity (STDP (Bi & Poo, 1998)); a more detailed version of the calcitron with temporal dynamics would be necessary to capture STDP.)

The fourth source of calcium, *C*_*sprv*_, comes from an external supervisor, denoted as *Z. Z* may be binary or positive real-valued. In hippocampal pyramidal neurons, a likely candidate for this supervisory signal is strong input to the apical tuft, which can induce bursts of spikes at the soma, potentially leading to global calcium influx at VGCCs at basal dendrites (Bittner et al., 2015, 2017a; Grienberger & Magee, 2022a; Milstein et al., 2021). A similar calcium-based supervisory scheme exists in cerebellar Purkinje neurons, where strong input to the Purkinje neurons from climbing fibers can induce long-term depression of synapses between presynaptic parallel fibers and the postsynaptic Purkinje cell (Konnerth et al., 1992). (In our model, *Z* does not contribute directly to the neuron’s output *ŷ*, because although the supervisory signal often does depolarize the neuron, this depolarization is treated as incidental to the plasticity induction, rather than as part of the neuron’s input-output function. In other words, we assume that downstream neurons only care about “output spikes” rather than “plasticity plateaus\bursts”). The calcium influx due to this supervisory signal is defined as:

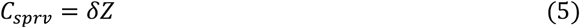

Where *δ* is the coefficient determining the amount of calcium that comes from the supervising signal. The total calcium per dendritic spine, 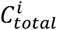, is the sum of these four calcium sources:

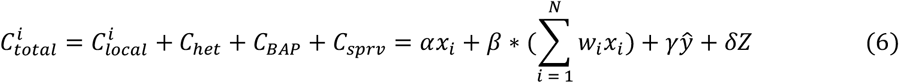

An equivalent way to write this equation is in terms of local and global calcium sources:

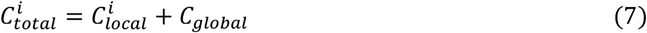

where:

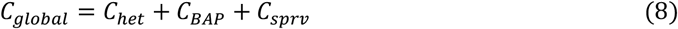

This formulation emphasizes that global calcium signals from the total feed-forward depolarization, backpropagating action potential, and supervisor are broadcast equally to all synapses, thus the local calcium is needed to break the symmetry between different synapses at each time step.

### Calcium-Based Plasticity for the Calcitron

At each time step *t* of the calcitron’s operation, a vector of inputs *x* = [*x*_1_, *x*_2_ … *x*_*N*_] is presented to the calcitron, and the calcitron produces an output *ŷ*(Eq. (1)). The calcitron calculates the calcium concentration per spine, 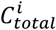, from the inputs (*x*), output (*ŷ*), and the supervising signal (*Z*) at that time step (Eq. (6)). The calcium concentration at each dendritic spine 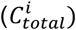 is used to determine the magnitude and direction of plastic change (if any) for that synapse’s weight (*w*_*i*_) at the next time step.

To implement calcium-based plastic changes to each synapse, we consider two versions of a calcium-based plasticity rule: a linear version and an asymptotic version. The latter rule is also referred to as the fixed point – learning rate (FPLR) rule. Although we will use the FPLR rule for most of the simulations in this study, we will first introduce the linear rule as it is simpler and provides a point of entry to the FPLR rule.

The linear version of calcium-based plasticity changes a synaptic weight by a fixed amount at each time step depending on the amount of calcium present at the synapse at that time step. Formally, following the notation of (Shouval et al., 2002) this may be described as

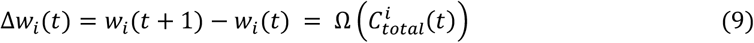

where *Ω*(*C*) is any function from positive-valued calcium concentrations to positive or negative changes in synaptic weight. As a simple representation of the classic calcium control paradigm observed in hippocampal and cortical cells, we can choose *Ω*(*C*) to be a step function with two thresholds: the depression threshold, *θ*_*D*_, and the potentiation threshold, *θ*_*P*_, where *θ*_*D*_ *< θ*_*P*_. Our plasticity function *Ω*(*C*) returns 0 when the calcium is below *θ*_*D*_, returns −0.01 when the calcium is between *θ*_*D*_ and *θ*_*P*_ (depressing the synapse by 0.01 units), and returns 0.1 when the calcium is above the *θ*_*P*_ (potentiating the synapse by 0.1 units). Formally this can be written as:

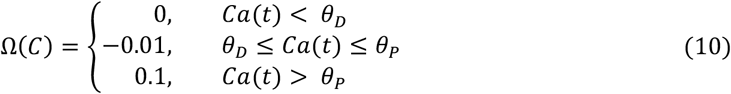

In the above formulation of the plasticity rule, synaptic weights can become arbitrarily large or small provided a sufficient number of plasticity events, including changing sign from positive (excitatory) to negative (inhibitory). This is not particularly biologically realistic, as synapses do not change sign from excitatory to inhibitory, and synaptic strengths are not observed to become arbitrarily large (there are also biophysical limitations on the maximum possible depolarization that can be achieved from a single synaptic input). We thus employ a different rule, inspired by the rules of Shouval and Graupner (Graupner & Brunel, 2012), wherein synaptic weights are modified asymptotically toward a minimum or maximum value (*w*_*min*_ and *w*_*max*_), depending on the calcium concentration. This is known as the fixed point – learning rate (FPLR) rule, as the rule is specified by defining the fixed points of the weights (i.e. the asymptotes, *w*_*min*_ and *w*_*max*_ – also called *F*_*D*_ and *F*_*P*_, respectively, as the depressive and potentiative fixed points) and learning rates (*η*_*D*_ and *η*_*P*_ for depression and potentiation, respectively) as a function of the calcium concentration (Moldwin, Azran, et al., 2023). In the absence of a substantial [Ca^2+^], i.e. (*Ca*(*t*) *< θ*_*D*_), the weight can also drift back to some baseline *w*_*drift*_ at a slower rate of *η*_*drift*_. The FPLR rule for the calcitron is formulated as:

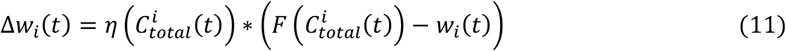

where *η*(*C*) defines the learning rate as a function of calcium and *F*(*C*) defines the fixed points as a function of calcium. The learning rates *η*(*C*) define the fraction of the difference between the present weight and the target fixed point that is traversed at each time step, resulting in asymptotic plasticity dynamics for a given level of calcium at a particular synapse. A standard two-threshold calcium control function would have the following structure:

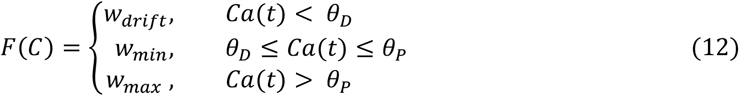

And

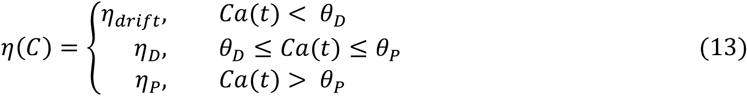

For example, we might have:

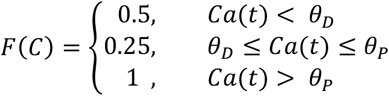

And

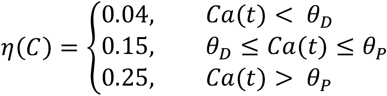

This means that synaptic weights with pre-depressive calcium concentrations 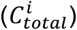 eventually drift toward a “neutral” state of 0.5 at a rate of 0.04 synapses with a depressive calcium concentration will depress towards 0.25 at a rate of 0.15, and synapses with a potentiative 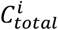 will be potentiated towards 1 at a rate of 0.25. In practice, for the purpose of this work, we will neglect the pre-depressive drift, i.e. the tendency of synapses to slowly drift back to baseline in the presence of very low levels of calcium (*C < θ*_*D*_). In the FPLR framework, this is accomplished by setting *η*(*C < θ*_*D*_) = 0.

### Calcitron leaning in a binary model: Hebbian, Anti-Hebbian, and other Pre-Post Learning Rules

As simple illustration of the ability of the Calcitron to implement learning rules, we consider Hebbian learning. One simple formulation of the Hebb rule is “neurons that fire together, wire together”. In other words, presynaptic inputs that were active at the same time as a postsynaptic spike are potentiated. In the Calcitron, if we assume binary inputs and outputs (i.e. *x*_*i*_ and *ŷ*are either 0 or 1), this can be accomplished by setting *α* and *γ* in Eq. (6) to both be below *θ*_*D*_, while enforcing that *α* + *γ* is above *θ*_*P*_. This ensures that an active synapse not accompanied by a spike does not change (because *α* * 1 + *γ* * 0 *< θ*_*D*_), nor does an inactive synapse that is accompanied by a spike (because *α* * 0 + *γ* * 1 *< θ*_*D*_), but an active synapse accompanied by a spike will potentiate (because *α* * 1 + *γ* * 1 > *θ*_*P*_) (Fig. 2A1).

**Figure 1:**
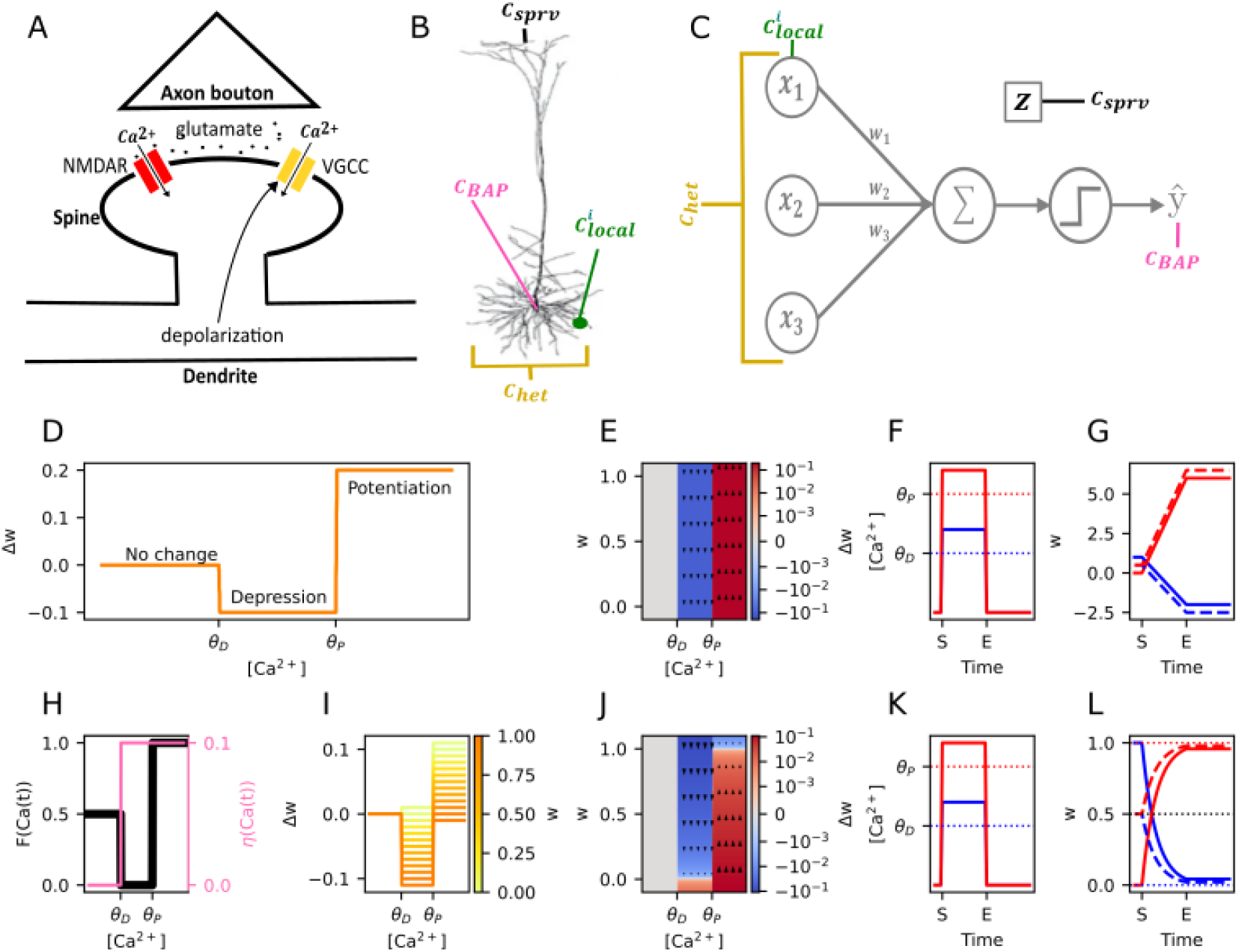
The Calcitron model and calcium-based plasticity rules. **(A)** Sources of Ca^2+^ at the synapse. Local glutamate release from an activated presynaptic axon binds to an NMDA receptor in the postsynaptic dendritic spine, enabling local Ca^2+^ influx. Depolarization of the neuron opens voltage-gated calcium channels (VGCCs), enabling calcium influx from global signals. (NMDA receptor voltage-dependence is neglected for simplicity). **(B)** Possible sources of Ca^2+^ influx in a neuron. Ca^2+^ can enter due to presynaptic input 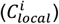, heterosynaptically-induced depolarization of VGCCs (*C*_*het*_), the backpropagating action potential (*C*_*BAP*_) or a supervisory signal, such as a somatic burst induced by input to the apical tuft (*C*_*sprv*_). **(C)** The four Ca^2+^ sources in a point neuron model. Each Ca^2+^ source is associated with a respective coefficient (*α, β, γ, δ*) determining how much Ca^2+^ comes from each source. **(D)** The calcium control hypothesis. [Ca^2+^] between *θ*_*D*_ and *θ*_*P*_ induces depression, [Ca^2+^] above *θ*_*P*_ induces potentiation. **(E)** Weight change as a function of calcium in the linear version of Ca^2+^-based plasticity, as in (D), shown as phase plane. Magnitude of weight change is independent of current weight. Blue indicates depression, red indicates potentiation, white indicates no change. **(F)** Step stimulus to show the plastic effect of different levels of [*Ca*^2+^]. [*Ca*^2+^] is either raised to a depressive level (*θ*_*D*_ ≤ [*Ca*^2+^] ≤ *θ*_*P*_, blue line) or to a potentiative level ([*Ca*^2+^] > *θ*_*P*_, red line) for several timesteps, then reduced to 0. S and E refer to start and end of stimulus. **(G)** Dynamics of the linear rule in response to the canonical stimulus from (F). Synaptic weights increase or decrease linearly in response to the potentiative or depressive levels of calcium (red and blue traces, respectively), then remain stable after calcium is turned off. **(H)** Fixed points (black) and learning rates (pink) in the asymptotic fixed point – learning rate (FPLR) version of the calcium control hypothesis. **(I)** Weight change as a function of [Ca^2+^] for different values of the current synaptic weight. Darker colors indicate higher weights. **(J)** Phase plane of weight changes for the FPLR rule. **(K)** Stimulus to demonstrate FPLR rule, identical to F. **(L)** Dynamics of the FPLR rule. Synaptic weights potentiate or depress asymptotically toward the potentiative or depressive fixed point.

**Figure 2:**
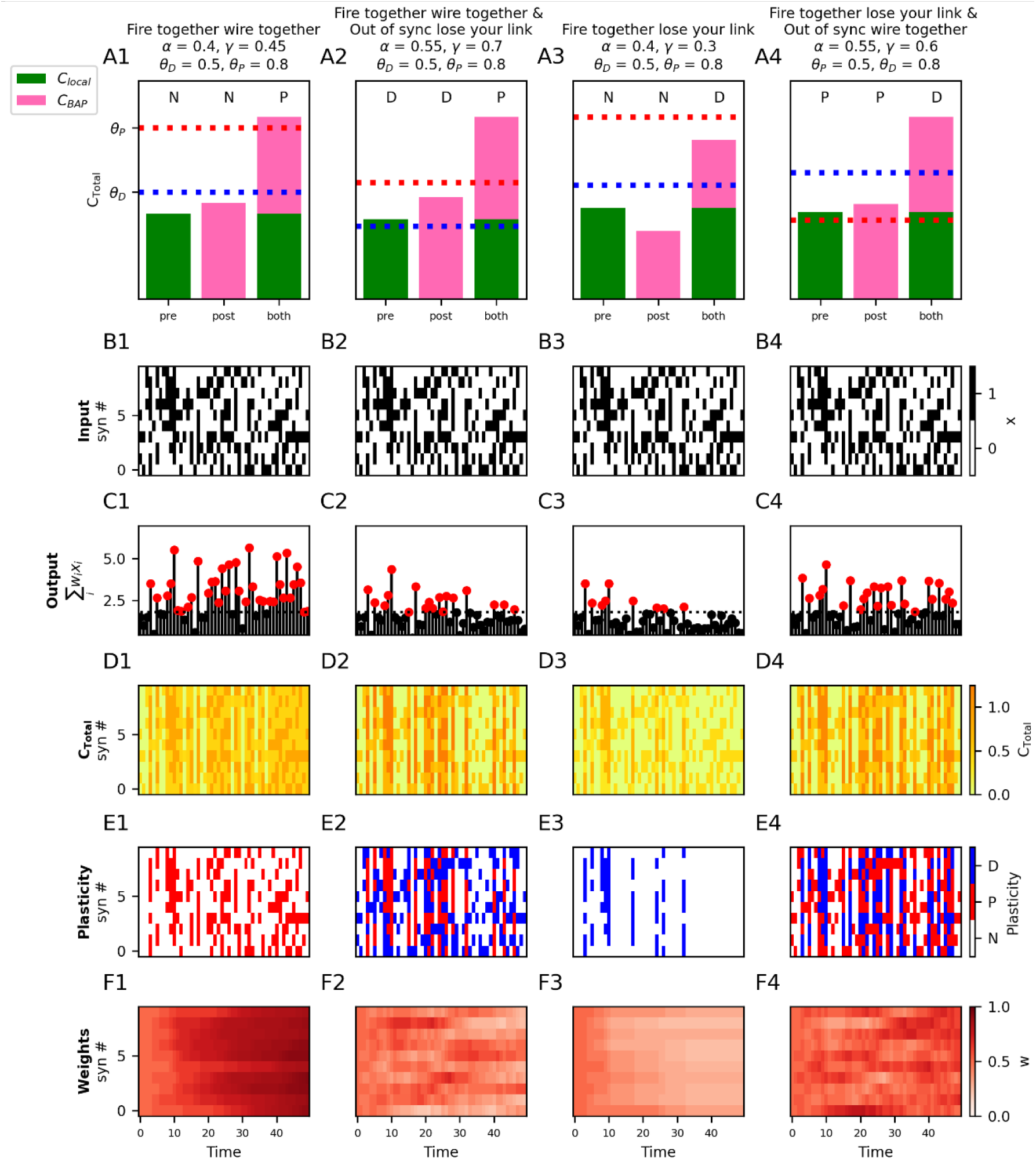
Four kinds of Hebbian and anti-Hebbian learning using Ca^2+^. **(A1-A4)** Different versions of Hebbian and anti-Hebbian learning rules are implemented by setting the respective coefficients (*α* and *γ* in Eq. 6) for the local 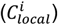 and backpropagating spike-dependent (*C*_*BAP*_) [Ca^2+^]. For each rule, we show the direction of plastic change (indicated by the letters above the bars: “N”: no change, “D”: depression, “P”: potentiation) for three different conditions: a synapse *with* active presynaptic input in the *absence* of a postsynaptic spike (“pre”), a synapse *without* local input in the *presence* of a postsynaptic spike (“post”) and at a synapse *with* active presynaptic input *and* a postsynaptic spike (“both”). The total [Ca^2+^] 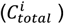 for each condition is the sum of the local input-dependent [Ca^2+^] (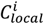, green) and the spike-dependent [Ca^2+^] (*C*_*BAP*_, pink). (When there is neither local input or a postsynaptic spike, the expected [Ca^2+^] is 0) (**B1 – B4)** For each learning rule from (A1-A4), 10 random binary inputs (black: active, white: inactive) are presented to each synapse at each time step. (Inputs are identical for all learning rules). **(C1-C4)** Sum of weighted inputs at each time step for each learning rule shown in A1-A4 respectively. Dotted horizontal line indicates the spike threshold (−*b* from equation 6). Outputs that are above the threshold (produce a postsynaptic spike) are indicated by a red circle. **(D1 – D4)** [Ca^2+^] per synapse for each time step for the 4 learning rules shown in A1-A4 respectively. **(E1 – E4)** Bar codes indicating occurrence of potentiation (“P”, red), depression (“D”) or no change (“N”, white) shown in A1-A4 respectively. **(F1-F4)** Synaptic weights over the course of the simulation for A1-A4 cases respectively.

Another version of the Hebbian learning rule can be stated as “fire together, wire together; out of sync, lose your link”. In other words, synapses that were active at the same time as a postsynaptic spike are potentiated, as before, but now we penalize (via depression) synapses that were active at a time when there was no postsynaptic spike, as well as synapses that were inactive at a time when a postsynaptic spike did occur. In the Calcitron, this is accomplished by setting *α* and *γ* to individually be in the depressive region (between *θ*_*D*_ and *θ*_*P*_), which penalizes ‘out of sync’ synapses, while still maintaining *α* + *γ* is above *θ*_*P*_ to enforce “fire together wire together” behavior (Fig. 2A2).

It is also possible to obtain an anti-Hebbian “fire together, lose your link” plasticity rule by setting *α* and *γ* to individually be below *θ*_*D*_, while enforcing that *θ*_*D*_ *< α* + *γ < θ*_*P*_. However, it is *not* possible, using the standard plasticity thresholds, to get an anti-Hebbian rule that rewards out-of-sync synapses, i.e. “fire together, lose your link, out of sync wire together”, as a synapse that fires synchronously with the postsynaptic spike will always have a higher 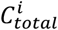 than a synapse that fires asynchronously, and thus cannot potentiate. In other words, if *θ*_*D*_ *< α* + *γ < θ*_*P*_, necessarily it holds that *α < θ*_*P*_ and *γ < θ*_*P*_ (Fig 2A3).

However, if the plasticity thresholds are reversed, i.e. *θ*_*P*_ *< θ*_*D*_, as in Purkinje neurons, we can get “fire together, lose your link; out of sync, wire together” plasticity. We set *α* and *γ* to both be in the potentiation region (between *θ*_*P*_ and *θ*_*D*_), while enforcing that *α* + *γ* is above *θ*_*D*_ (Fig. 2A4).

As a simple demonstration of these Hebbian and anti-Hebbian rules, we created a random sequence of binary input patterns (Fig. 2B1-2B4) consisting of 10 synaptic input lines (i.e., N=10) and presented this sequence to four different Calcitrons, each implementing one of four rules described above by applying different calcium source coefficients and plasticity thresholds. We then compare how the different Calcitrons yield different weight dynamics (Fig. 2C-2F).

In the “fire together, wire together” Calcitron (2A1-2F1), synaptic weights can only potentiate or stay stable, they can never depress. Every so often, just by chance, one of the random input patterns will be sufficiently large to generate a spike (2C1, red-headed stems). The calcium from this spike combined with the calcium at active synapses yields potentiation at the active synapses (2D1-2E1). This creates a positive feedback loop: when a synapse is potentiated, that makes the neuron more likely to elicit a spike whenever that synapse participates in a pattern (because they synapse provides a greater contribution to the overall input to the neuron). Eventually, this results in all synaptic weights becoming strongly potentiated (2F1) and the Calcitron spikes in response to almost all input patterns.

In the “fire together, wire together; out of sync, lose your link” Calcitron (2A2-2F2), the potentiation at active synapses whenever there is a spike is counterbalanced by depression at inactive synapses when a spike occurs, as well as at active synapses when no spike occurs. As such, the synaptic weights do not become overly strong and the neuron doesn’t spike as aggressively.

In the “fire together, lose your link” Calcitron (2A3-2F3), synaptic weights can only depress or stay stable, they can never potentiate. As such, initially, whenever the random input elicits a spike, the synapses that were active at that time step depress. However, here we have a negative feedback loop: once a synapse depresses, it contributes less to the neuron’s overall voltage, which makes the neuron less likely to spike, so eventually the synaptic weights stop depressing.

Finally, in the “fire together, lose your link; out of sync, wire together” Calcitron (2A4-2F4), we again have a balance between depression and potentiation for synchronized and unsynchronized activity, so we again observe more moderate changes in the synaptic weights over longer time horizons.

Importantly, it is possible to implement many more pre-post rules with calcium than just the standard Hebbian and anti-Hebbian rules. For example, we could have a “fire together wire together” rule that penalizes synapses that are active when there is no postsynaptic spike, but doesn’t penalize inactive synapses at the time when there is a postsynaptic spike (Figure 3, third row first column).

**Figure 3:**
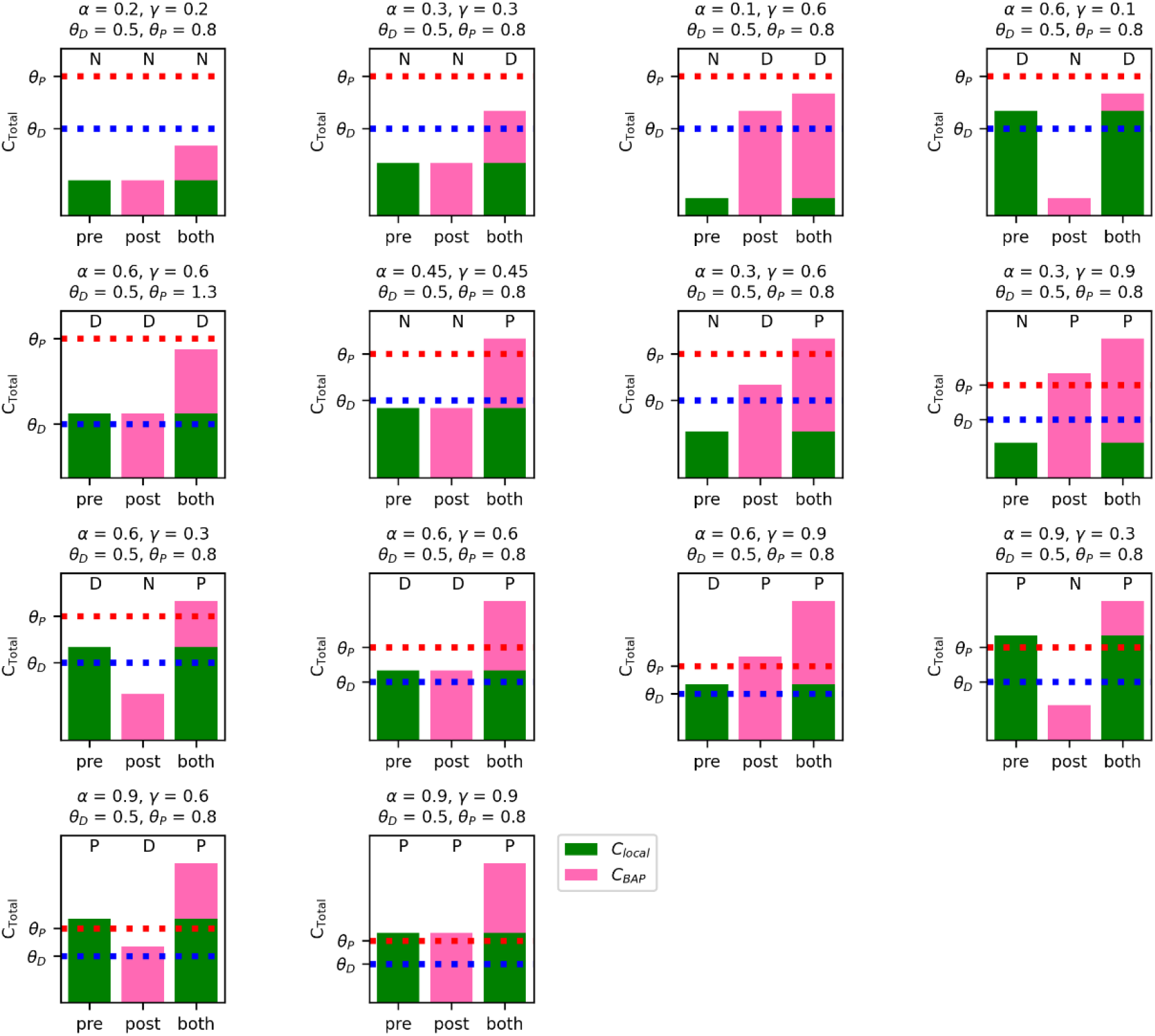
Fourteen possible plasticity rules for presynaptic input- and spike-dependent-calcium. Setting different coefficient values (*α* and *γ* in Eq. 6) for the local 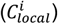 and backpropagating spike-dependent (*C*_*BAP*_) [Ca^2+^] can lead to fourteen possible learning rules in the case of binary input and output. Presynaptic input alone (standalone green bar in each subplot) can lead to potentiation (P), depression (D), or no change (N) at the active synapse, or a postsynaptic spike alone (standalone pink bar in each subplot) can lead to P,D, or N at non-activated synapses. When a postsynaptic spike occurs at the same time as a presynaptic input (stacked pink and green bars in each subplot), the resultant [Ca^2+^] at an activated synapse is the sum of the local and spike-dependent [Ca^2+^], constraining the number of qualitatively different rules to the 14 shown here (assuming *θ*_*P*_ > *θ*_*D*_). Combinatorically, there are 27 possible rules (every combination of {P, D, N} with {pre, post, and both}, however only the 14 listed are consistent with the calcium control hypotheses.)

From a combinatoric standpoint, one can imagine 27 possible pre-post learning rules, because there are three possible synaptic scenarios (pre only, post only, pre and post) and there are three possible outcomes for each case (potentiate, depress, nothing). (If we allow for synapses to undergo plasticity when there is neither presynaptic nor postsynaptic activity, there are 81 possibilities, but we assume that synapses are stable in the absence of any activity.) However, the two-threshold calcium control hypothesis prohibits some of these scenarios, because the [Ca^2+^] from the “pre and post” scenario most be the sum of the [Ca^2+^] from the “pre only” and “post only” scenarios. This constraint leaves us with 14 potential pre-post rules that can be implemented in the calcitron (Fig. 3).

### Frequency-dependent pre-post learning rules in a rate model

In the previous section, we assumed binary inputs and outputs, (i.e. *x*_*i*_ and *ŷ*are either 0 or 1). If we instead consider a rate model, where *x*_*i*_ and *ŷ*can be any positive value, it is possible to implement frequency-dependent plasticity whose outcome depends on both the pre-synaptic and postsynaptic firing rate (Fig. 4A). Experimentally, we note that presynaptic-only frequency-dependent plasticity has been observed, where low frequency stimulation causes depression and high frequency stimulation causes potentiation (Bliss & Lomo, 1973; Dudek & Bear, 1992; O’Connor et al., 2005a). To replicate classical (presynaptic only) frequency-dependent plasticity, we set all coefficients other than *α* to 0. The direction of plasticity at any given synapse will depend on the input strength, i.e. the value of *x*. According to Eqs. 6 and 12, if *x* is between 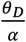 and 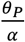, *w*_*i*_ potentiates, if *x* is larger than 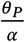, *w*_*i*_ depresses, otherwise there is no change (Fig. 4B1).

**Figure 4:**
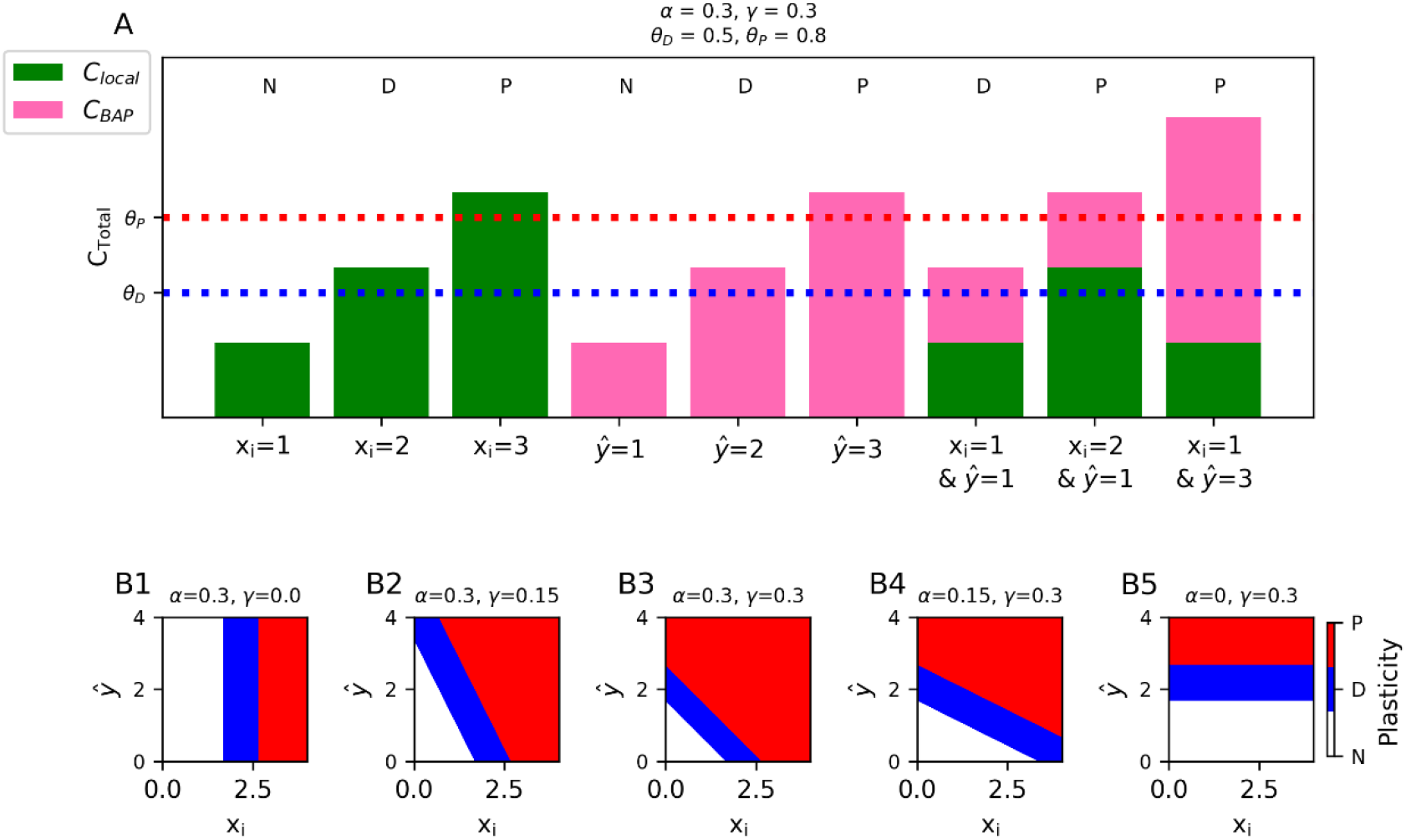
Frequency-dependent pre- and post-synaptic plasticity in rate-based models. **(A)** Calcium-dependent plasticity in a rate model. In the absence of postsynaptic spikes, sufficiently strong presynaptic inputs (x_i_) alone can generate plasticity at active synapses (green bars). Postsynaptic firing alone (y_i_) can induce plasticity even at inactive synapses (pink bars), and the combination of presynaptic input and postsynaptic spiking can sum to induce plasticity at active synapses (stacked green and pick bars). **(B1-B5)** [Ca^2+^] (binned into regions of no change (white), depressive (blue) or potentiative (red)) as a function of presynaptic (x-axis) and postsynaptic (y-axis) firing rate. Each panel has a different value for *α* ([Ca^2+^] per presynaptic spike) and *γ* ([Ca^2+^] per postsynaptic spike).

If we choose nonzero values for both *α* and *γ*, the direction of plasticity at each synapse will depend on sum of the strength of its presynaptic input *x*_*i*_ and well as the output strength *ŷ*. By choosing different values for *α* and *γ*, it is possible to emphasize the effect of the presynaptic input versus the postsynaptic output on the [Ca^2+^], and consequently the plasticity (Figure 4B2-4B4).

It is similarly possible to have postsynaptic-only frequency-dependent plasticity, which depends only on the output strength, *ŷ*, by setting all coefficients other than *γ* to 0. Now, the plasticity at all synapses will depend only on the output firing rate,*ŷ*. If *ŷ*is between 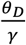 and 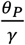, all synapses potentiate, if *ŷ*is larger than 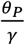, all synapses depress, otherwise no change occurs at any synapse (Fig. 4B5).

Because we are using a step function for the learning rate *η*(*C*) (Eq. (11)), the frequency of the input and output affect only the direction of the synaptic change, not its magnitude. However, if desired, it is possible to implement more biologically realistic frequency-dependent rules where the magnitude of plasticity is more precisely titrated by the [Ca^2+^] by defining *η*(*C*) as a soft threshold function (e.g., a sum of sigmoid functions) instead of a step function (see (Moldwin et al., 2023)).

### Unsupervised learning of repetitive patterns with heterosynaptic plasticity

One task we might want a neuron to perform is to learn recognize a particular input pattern, i.e., by emitting a spike in response to it while not firing in response to other input patterns. This task is usually implemented as a supervised learning task (such as in the perceptron algorithm, see below), but it is also possible for a neuron to learn to recognize specific patterns in an unsupervised fashion using heterosynaptic plasticity. Instead of directly telling the neuron which inputs should elicit a spike, it is possible to teach a neuron using heterosynaptic plasticity to spike only in response to frequently repeated “signal” patterns, while ignoring sporadic random “noise” patterns (Fig. 5A).

**Figure 5:**
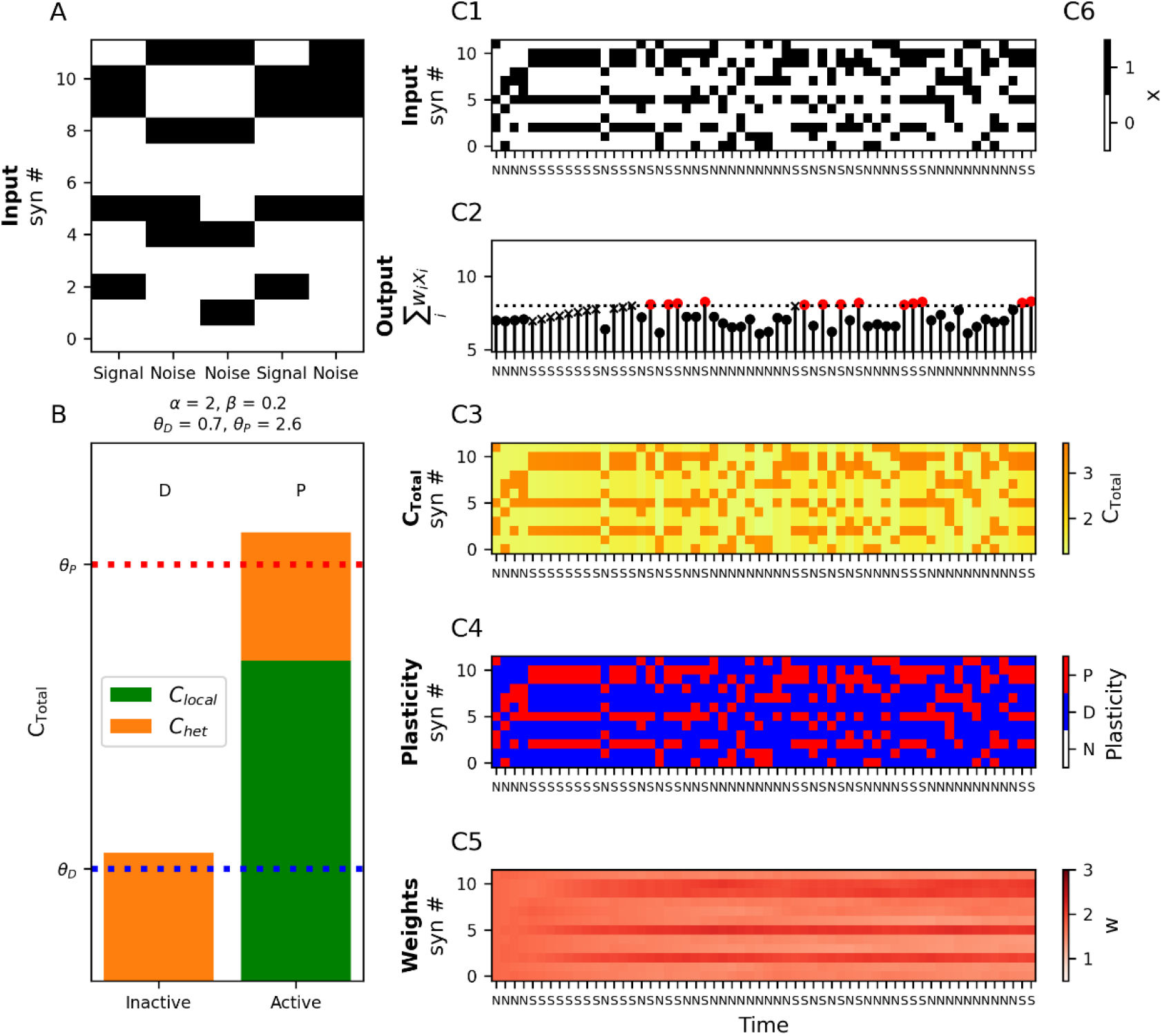
Learning to recognize repetitive patterns with heterosynaptic plasticity. **(A)** A “signal” pattern is presented repeatedly to the neuron interspersed with non-repeating random “noise” patterns of the same sparsity. **(B)** Within each input pattern (both signal or noise) inactive synapses depress (above *θ*_*D*_ at left) due to the heterosynaptic calcium, whereas active synapses will potentiate from the sum of heterosynaptic calcium, *C_het_*, and local calcium *C_local_* (above *θ_P_* at right). **(C1)** Signal and noise patterns are presented to the neuron. (S: signal, N: noise). **(C2)** Spiking output of the calcitron. Black: no spike, Red: spike. An ‘x’ marker indicates incorrect output (e.g. no spike in response to a signal pattern), filled circles indicate correct outputs. Note the increase in correct spiking output for the signal inputs depicted by the red circles. **(C3-C5)** Calcium, plasticity, and weights over time respectively as the input patterns in C1 are presented. Note that the synaptic weights (C5) change so that they eventually resemble the signal patterns.

We assume here that both the signal and noise input patterns here have the same sparsity, i.e. that there are always *k* out of *N* active synapses at every time step. We enforce that input patterns always heterosynaptically depress non-active synapses by setting *β* in Eq. 6 such that *θ*_*D*_ *< βkw*_*min*_ *< βkw*_*max*_ *< θ*_*P*_ and homosynaptically potentiate active synapses by enforcing that *βkw*_*min*_ + *α* > *θ*_*P*_ (Fig. 5B).

Every time an input pattern is presented to the neuron, the synapses that are active in that pattern will be potentiated and the synapses that are inactive will be depressed. If *η*(*C*) *<* 1, the potentiation and depression will occur gradually as the patterns are presented, and each synapse thus retains a “memory” of the recent input history. This creates a sort of competition between input patterns, because each pattern potentiates its active synapses while depressing synapses that were active in other patterns. Patterns that are repeated frequently (the signal patterns) will tend to dominate this competition, as the other (noise) non-repeated patterns will cancel out each other’s plastic influence on the synaptic weights. Over time, the calcitron’s synapses will tend to strengthen the synapses associated with the frequently repeated signal pattern and depress other synapses, eventually inducing the calcitron to spike only in response to the signal pattern (Fig. 5C1-C5).

Very loosely speaking, the above plasticity rule can be understood as a process to set the neuron’s weights to a running average over its recent inputs. This is why the neuron is able to detect the signal patterns from the noise patterns – because the noise patterns average each other out, leaving only the active synapses from the signal pattern. Because of this averaging effect, if there were multiple repeating input patterns, we would expect the neuron’s weights to be strongest at synapses that participate in more patterns and weaker at synapses that only participate in a few patterns. We note that because this learning rule only depends on the weighted inputs and not the activation function *g*, the learning rule is similar in structure and function to Oja’s rule for neural learning of principle components (Oja, 1982). For additional work on plasticity models for unsupervised learning of input patterns, see (Clopath et al., 2010; Yeung et al., 2004).

### “One-shot flip-flop” (1SFF) model of behavioral time-scale plasticity (BTSP)

Recent experimental findings in the hippocampus have revealed a novel form of plasticity, known as behavioral time scale plasticity (BTSP) (Bittner et al., 2015, 2017b; Grienberger & Magee, 2022b; Milstein et al., 2021). A mouse running on a treadmill can spontaneously form a place field in a CA1 hippocampal neuron when the soma of the neuron in injected with a strong current, inducing a plateau potential (this also occurs spontaneously in vivo via a supervising signal from the entorhinal cortex, see (Bittner et al., 2017; Grienberger & Magee, 2022). After a single induction, this plateau potential results in the neuron exhibiting a place field selective to the mouse’s location few seconds before or after the time of the plateau potential. Moreover, this place field can be modified; if a second plateau potential is induced while the mouse is at a different location near the first place field, the place field will shift to the new location, thus “overwriting” the first place field (Milstein et al., 2021). In previous work, we showed how the calcium control hypothesis might be able to explain various aspects of these experimental results (Moldwin et al., 2023).

BTSP can be reduced to a more abstract, idealized form of learning. If we ignore the precise temporal dynamics, BTSP can be thought of as a form of supervised one-shot learning, wherein a supervisory signal potentiates all synapses that were coactive with it, while depressing all other synapses. If we consider a bistable synaptic weight that can either be in an “potentiated” (1, i.e., *w*_*max*_) or “depressed” (0, i.e., *w*_*min*_) position, this supervisory plateau signal in BTSP is effectively a “write” command which tells the neuron to store the state of its inputs, i.e. active (1) or inactive (0) as its weights. We call this “one-shot flip-flop” learning, because the supervising signal overwrites all the weight to new binary values in a single timestep.

To implement this in the calcitron, we again assume binary inputs, and we also enforce binary synaptic weights by setting *η*(*C*) = 1 in both the depressive and potentiative regions of calcium, so a synapse will immediately be set to *w*_*min*_ whenever the [Ca^2+^] in the depressive region and to *w*_*max*_ whenever there is a potentiative [Ca^2+^] value (Fig. 6A1-A3).

**Figure 6:**
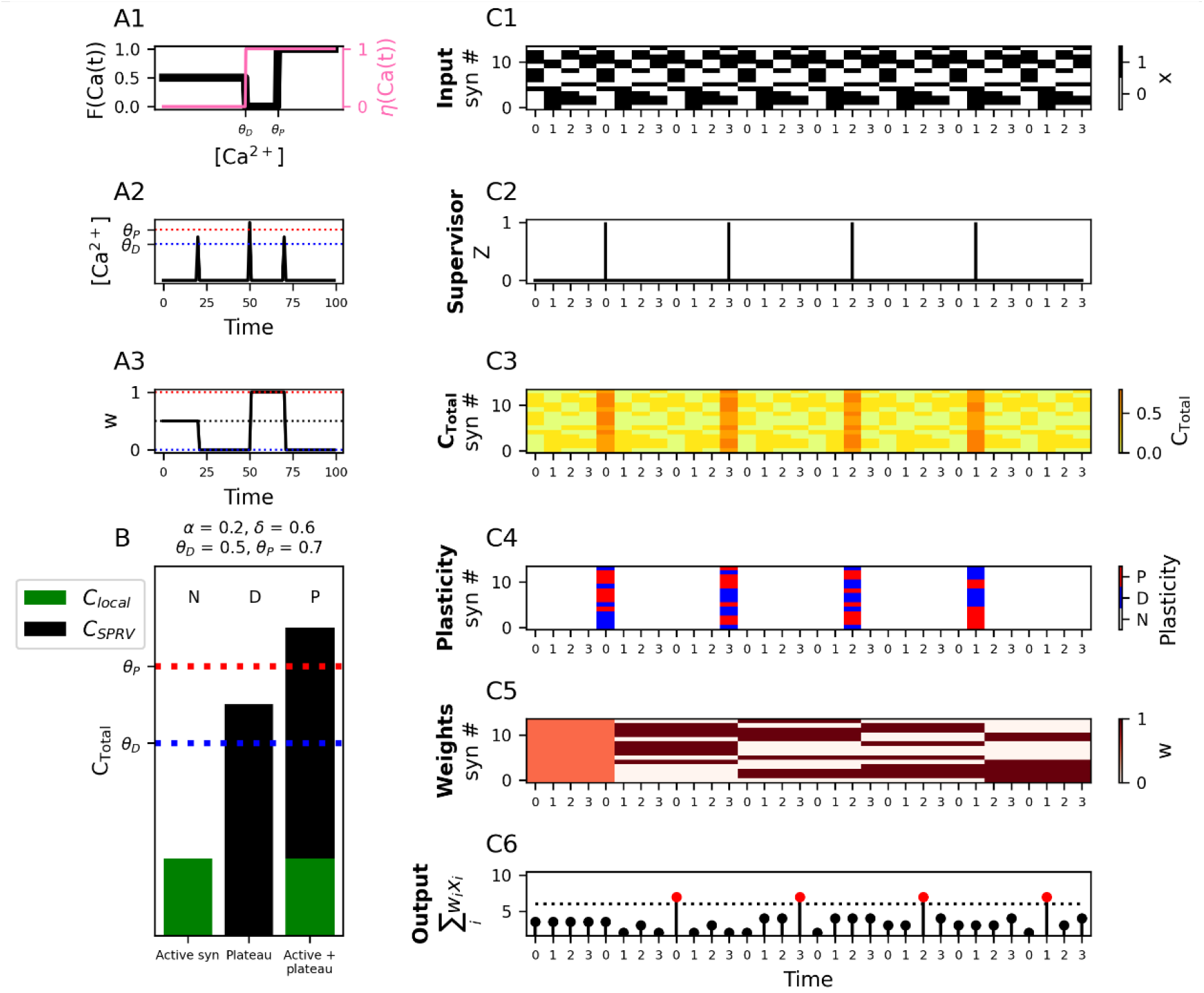
“One-shot flip-flop” (1SFF) plasticity. **(A1)** Fixed points (*F*(*Ca*(*t*)), black line, left y-axis) and learning rates (*η*(*Ca*(*t*)), pink line, right y-axis) for the different regions of [*Ca*^2+^]. For 1SFF learning, the learning rate is set to 1 in the depressive and potentiative regions of [*Ca*^2+^] for immediate switch-like plasticity. **(A2)** Exemplar stimulus illustrating plasticity dynamics. An instantaneous [*Ca*^2+^] pulse is generated at three timesteps over the course of the experiment. **(A3)** Synaptic weights over time in response to stimulus presented in A2. **(B)** 1SFF plasticity rule. Local input alone does not reach the depression threshold, a plateau potential alone induces a depressive [*Ca*^2+^], but local input combined with a plateau potential induces potentiation. **(C1)** A repeated sequence of input patterns (0,1,2,3) corresponding to locations on a circular track that a mouse traverses at each timestep as it runs multiple laps. **(C2)** Externally generated supervisory signal (plateau potential), Z, presented at different locations over the course of the experiment. **(C3-C5)** [*Ca*^2+^], plasticity, and weights, respectively for each time step. Synaptic inputs at the time of the supervisory signal are “written” to the synaptic weights at the following respective time step. **(C6)** Neural output at each time step. Red circles indicate spikes. Note that the neuron spikes at the location at which the supervisory signal occurred in the previous lap.

Assuming that the supervisory signal *Z* = 1 when active, one-shot flip-flop learning can implemented in the calcitron by enforcing *α < θ*_*D*_ (presynaptic input alone doesn’t cause plasticity), *θ*_*D*_ *<* δ *< θ*_*P*_ (the supervisory signal depresses all inactive synapses) and *δ* + α > *θ*_*P*_ (the supervisory signal potentiates all active synapses) (Fig. 6B).

To illustrate the effect of one-shot flip-flop learning, we present the calcitron with a repeated sequence of 4 binary input patterns, labeled with the numbers 0-3, to simulate a mouse running on a circular track (Figure 6C1). These 4 inputs patterns can be thought of as locations on a track traversed by the mouse. We then provide a supervisory provide a supervisory signal at random intervals (Figure 6C2). Every time the supervisory signal is presented, the binary state of the weights at the following time step are set to the binary state of the inputs that were active in tandem with the supervisory signal (Fig. 6C3-C5). The next time the neuron encounters the input patter (location) at which it previously received a supervisory signal, the neuron fires. In other words, the supervisory signal turned the neuron to a place field for that location. When another supervisory signal comes at a different location, the previous place field is overwritten with a new place field at the new location (Fig. 6C6).

### Homeostatic Plasticity with both Internal and Circuit Mechanisms

Another important form of plasticity does not involve storing new information per se, but rather maintaining a regular average firing rate. This form of plasticity is known as homeostatic plasticity. Homeostatic plasticity can be important for the health of the neuron (i.e. too much firing can deplete neuronal resources, potentially leading to cell death) as well as maintaining stability and regularity within neural circuits (G. Turrigiano, 2012; G. G. Turrigiano, 2008; G. G. Turrigiano et al., 1998; G. G. Turrigiano & Nelson, 2004).

Before we demonstrate how homeostatic plasticity can be implemented in the calcitron, we first note that our solution for the mechanism of homeostatic plasticity differs from what has been observed experimentally, which involves a different calcium signaling pathway and takes place over much longer time scales (many hours, instead of seconds or minutes) (G. Turrigiano, 2012). One form of experimentally-observed homeostatic plasticity seems to depend on *somatic* calcium concentrations – when a neuron is firing too slowly, the low calcium levels set off a signaling pathway to globally increase synaptic strengths, and when a neuron fires too much, the high calcium levels initiate a signaling pathway to decrease synaptic strengths. This form of homeostatic plasticity thus involves a different relationship between calcium and the direction of plasticity – low levels of somatic calcium induce potentiation, medium levels induce no change, and high levels induce depression (G. Turrigiano, 2012). For the purposes of our work, however, we propose an alternative strategy that depends on the calcium at the spine with the standard plasticity thresholds: low levels of calcium induce no change, medium levels induce depression, and high levels induce potentiation. As such, the calcitron version of homeostatic plasticity is a speculative exploration of how the brain could potentially implement homeostatic plasticity using the standard synaptic plasticity mechanisms and calcium thresholds.

In a rate model version of the calcitron, we can think of homeostatic plasticity as the problem of trying to keep the output firing rate, *ŷ*, within a target range, between 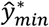 and 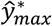. We will also assume that due to the postsynaptic refractory period, neurons have a maximum rate that it is physically possible to fire at, *ŷ*_*max*_, and that neurons also have a minimum physically possible firing rate *ŷ*_*min*_ (trivially, neural firing rates can’t be negative, so we will always have *ŷ*_*min*_ ≥ 0). In general, then, we have that 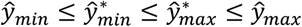.

For simplicity, if we assume that the synaptic inputs to the calcitron are binary (even though the output is not), there are broadly two strategies we can take with a homeostatic plasticity rule. One strategy, which we term “global homeostasis”, is that whenever the calcitron’s output *ŷ*is too large (or too small), we can depress (potentiate) all the calcitron’s synapses irrespective of their input. This will eventually result in the calcitron’s output being within the target range on average.

Although global homeostasis may be effective if the calcitron’s output is consistently too low or too high irrespective of the input patterns, globally depressing or potentiating all synapses is a drastic measure that can destroy previously stored information (unless scaling is multiplicative, see (G. G. Turrigiano, 2008; G. G. Turrigiano & Nelson, 2004)). Instead, we can use a more fine-grained approach, “targeted homeostasis”, which only modifies the synapses that were active at the time when the output firing rate was out of range, thus only correcting “errant” synapses. (See also (Rabinowitch & Segev, 2006b, 2006a) regarding the consequences of the global vs. local strategies of homeostatic plasticity when considering a realistic dendritic tree.)

For the global homeostasis strategy we will always set α = 0 so we can ignore presynaptic calcium, and for the targeted homeostasis strategy we will always set 0 *<* α *< θ*_*D*_ so the presynaptic calcium can break the symmetry between active and inactive synapses but isn’t so large that it is sufficient on its own to induce plasticity.

We first consider how both homeostatic plasticity strategies might be implemented using internal mechanisms (i.e. without an external supervisory signal). If our neuron is firing too strongly, 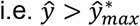, we would like to be the resultant calcium to be in the depressive [Ca^2+^] region. For the global homeostasis strategy, this can be implemented by setting 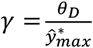 and *θ*_*P*_ > *γŷ*_*max*_, and in the targeted homeostasis strategy by setting 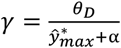 and *θ* > *γŷ*_*max*_ + α. Because *θ*_*P*_ > *γŷ*_*max*_ in both strategies, there is no longer any way to potentiate synapses, because even the maximum physically possible firing rate will not produce enough [Ca^2+^] to induce potentiation. We must instead rely on an external supervisory mechanism to potentiate synapses when the calcitron’s output is too low.

To construct a potentiation supervisor, we consider a simple disinhibitory circuit. The calcitron forms a synapse onto an inhibitory neuron, which inhibits a supervisory neuron that supervises the calcitron. When the calcitron’s output rises above 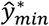, the inhibitory neuron is active, thus preventing the supervisory neuron from sending the calcitron an inhibitory signal. When the calcitron’s output falls below 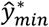, however, the inhibitory neuron becomes inactive, thus permitting the supervisory neuron to send a potentiative supervisory signal *Z*_*P*_ = 1 to the calcitron (Fig. 7A-7B). For the global homeostasis strategy, by setting δ = *θ*_*P*_ + ϵ, the supervisor induces potentiation at all synapses whenever 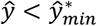. For the targeted homeostasis strategy, we set δ = *θ*_*P*_ − α + ϵ to ensure that only active synapses are potentiated, and we apply an additional constraint that δ *<* θ_D_, to ensure that non-active synapses aren’t depressed by the supervisory signal (Fig. 7C). (This constraint imposes that α *< θ*_*P*_ − θ_*D*_, which in turn entails that the pre-depressive region of [Ca^2+^] is larger than the depressive region of [Ca^2+^].)

**Figure 7:**
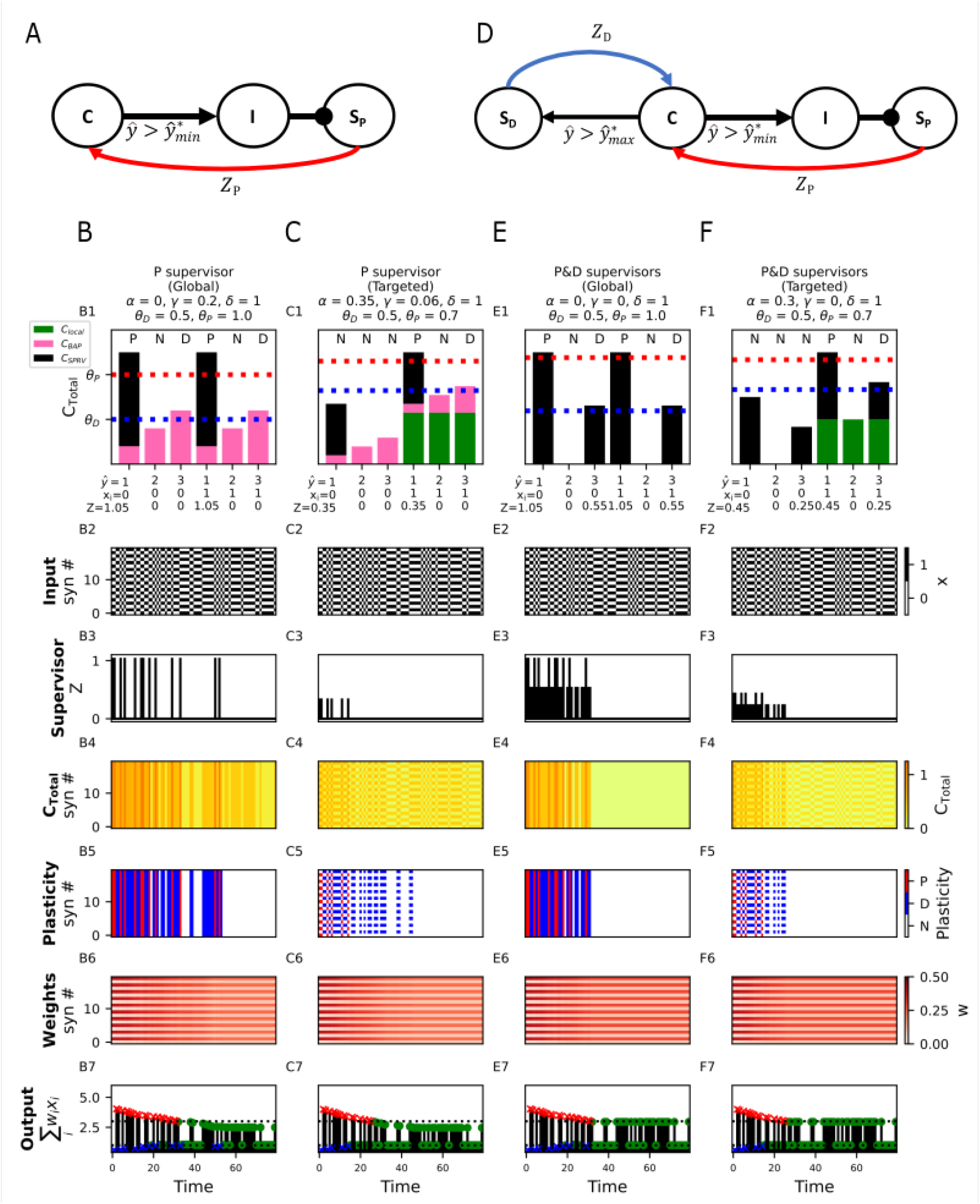
Different mechanisms for Homeostatic Plasticity. **(A)** Supervision circuit for homeostatic plasticity using only a potentiation supervisor. If the calcitron (“C”) fires above the target minimum rate (i.e.,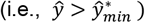) it activates an inhibitory population (“I”), which prevents the potentiation supervisor (*S*_*P*_) from producing a supervisory signal (in this case, *Z*_*P*_ = 0). If the calcitron’s output falls below its target range 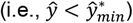 the supervisor is disinhibited, sending a potentiative calcium signal (*Z*_*P*_) to the calcitron. **(B)** Plasticity rule for global homeostatic plasticity using an internal mechanism for depression and a circuit mechanism for potentiation, as in (A). Here 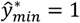 and 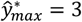. **(B1)** Overly strong outputs (*ŷ*> 3) produce sufficient calcium to depress all synapses; Overly weak outputs (*ŷ<* 1) result in the activation of *S*_*p*_, setting *Z*_*P*_ = 1, potentiating all synapses. **(B2)** Two input patterns (only even synapses or only odd synapses) are presented to the neuron in random order. **(B3)** *Z*_*P*_ occurs whenever *ŷ<* 1. **(B4-B6)** [Ca^2+^], plasticity, and weights over the course of the simulation. Even-numbered synapses are initialized to low weights; odd-numbered synapses are initialized to large weights. **(B7)** Neural rate output. Blue ‘x’ indicates output below 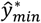, (lower dashed line), red ‘x’ indicates output above 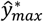 (upper dashed line), green circles indicate output in the acceptable range. **(C)** Plasticity rule for targeted homeostatic plasticity using the circuit from (A) as well as local [Ca^2+^]. **(D)** Supervision circuit for homeostatic plasticity using both a potentiation (*S*_*P*_) and depression (*S*_*D*_) supervisor. In addition to the disinhibitory circuit for the control of *S*_*P*_ as in (A), when the calcitron’s output is above the target output range 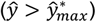, a depression supervisor (*S*) is activated, sending a depressive signal (*Z*) to the calcitron. **(E)** Plasticity rule for global homeostatic plasticity using both *S*_*P*_ and *S*_*D*_. **(F)** Plasticity rule for targeted homeostatic plasticity using *S*_*P*_ and *S*_*D*_ in combination with local calcium at active synapses.

It is also possible to implement both homeostatic potentiation and depression using external supervisors, instead of using an internal supervisor for depression. To do this, we set *γ*=0, so the postsynaptic spike itself doesn’t induce calcium influx, and we use the same disinhibitory circuit mechanism for depression as described above. To implement an external depression supervisor, we consider an addition circuit mechanism where the calcitron also synapses directly onto a new “depression supervisor” neuron, which synapses back onto the calcitron with a supervising synapse. This depression supervisor will be active whenever the calcitron’s firing rate *ŷ*exceeds *ŷ*_*max*_ (Fig. 7D).

Importantly, the depression supervisor and potentiation supervisor give supervisory signals of different strengths, *Z*_*D*_ and *Z*_*P*_, respectively. Without loss of generality, we can set *γ* = 1, so it is only necessary to set magnitude of the supervising signals. For the global homeostasis strategy, we set *Z*_*D*_ = θ_D_ and *Z*_*P*_ = θ_P_ + ϵ, which provides the calcitron with a global potentiative signal when 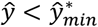 and a global depressive signal when 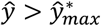 (Fig. 6E). For targeted homeostasis, we set *Z*_*D*_ = θ_D_ + α and *Z*_*P*_ = θ_P_ + α + ϵ, and we enforce that *Z*_*D*_ *< Z*_*P*_ *<* θ_D_ so that the supervising signals only modify active synapses (Fig. 6F).

To compare the two different supervisory circuits (external potentiation and internal depression vs. external potentiation and depression) and the targeted vs. global strategies, we created a calcitron whose even-numbered synapses were initialized to small weights and whose odd-numbered synapses were initialized to large weights. We then randomly present input patterns that either only activate the even-numbered synapses (“even patterns”) or only activate the odd-numbered synapses (“odd patterns). Initially, the even patterns produce an output which is too low (i.e., below 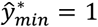) and the odd patterns produce an output which is too high (i.e., above 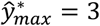). Over the course of presenting the patterns multiple times, the homeostatic mechanisms succeeded to increase the weights for the even synapses and decrease the weights for the odd synapses such that eventually the neuron fired within the target output range 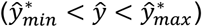 in response to both even and odd patterns. All four combinations of supervisory circuit structure and plasticity strategy (global vs. targeted) succeeded in this task, demonstrating that the same desired computational result can emerge from different underlying mechanisms (Fig. 6).

### Perceptron learning algorithm with calcium-based plasticity

We now show that it is possible to implement the perceptron learning algorithm with calcium-based plasticity. The perceptron learning algorithm (Rosenblatt, 1958) is the procedure by which a single linear neuron (see Eq. (1)) can learn to solve classification tasks, such as distinguishing between images of cats and dogs, by modifying its synaptic weights.

The perceptron learning rule is a supervised learning rule. Namely, each input pattern comes with an associated target outcome – that the neuron should either spike or not spike. Formally, we have a set of *P* input patterns {*x*^1^ … *x*^*μ*^ … *x*^*P*^} and associated labels {*y*^1^ … *y*^*μ*^ … *y*^*P*^}, where the bolded *x*^*μ*^ is an *N***-**dimensional vector of activity of the *μ*th input pattern and *y*^*μ*^ *∈* {0,1} is the associated target label, or class. The goal of the perceptron learning algorithm is to ensure that the neuron’s output on each pattern (*ŷ*^*μ*^) matches the target label, i.e., *ŷ*^*μ*^ = *y*^*μ*^.

The perceptron rule states that if the neuron makes an error on an input pattern *ŷ*^*μ*^ ≠ *y*^*μ*^, we increase or decrease the perceptron’s synaptic weights in a manner proportional to the input vector. Formally, if *ŷ*^*μ*^ = 0 and *y*^*μ*^ = 1 (a false negative; the neuron should have spiked but didn’t) we update each weight *w*_*i*_ according to the rule *w*_*i*_ ← *w*_*i*_ + η*x*_*i*_, where η is the learning rate. If *ŷ*^*μ*^ = 1 and *y*^*μ*^ = 0 (a false positive; the neuron spiked when it wasn’t supposed to) we update each weight *w*_*i*_ according to the rule *w*_*i*_ ← *w*_*i*_ − η*x*_*i*_. If the neuron produced the correct output, *ŷ*^*μ*^ = *y*^*μ*^, we don’t modify the weights. The changed in each synaptic weight at each time step, Δ*w*_*i*_(*t*) = *w*_*i*_(*t* + 1) − *w*_*i*_(*t*), for the perceptron learning rule can be described by Table 1.

**Table 1:**
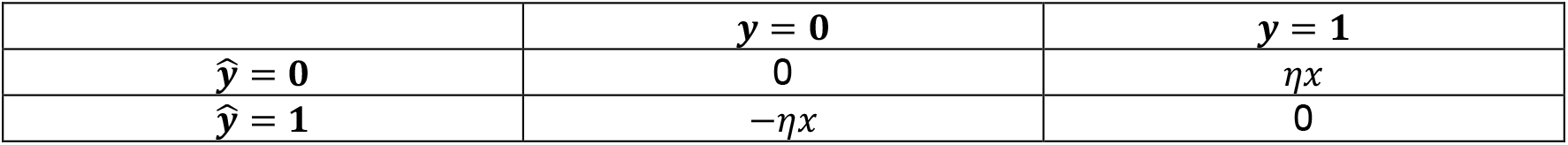
Weight update Δw in the standard perceptron learning rule.

We first note that because the fixed-point learning rate (FPLR) framework for calcium-based plasticity has asymptotic, weight-dependent dynamics, it is only possible to exactly replicate the perceptron learning rule with the linear calcium-based rule (Eq. (9)). However, because we would like to demonstrate perceptron-like dynamics with the more realistic FPLR rule (Eq. (11)), we propose an “asymptotic perceptron learning rule” which functions similarly to the original perceptron rule, except that weights increase or decrease asymptotically towards *w*_*min*_ or *w*_*max*_ instead of being able to increase or decrease indefinitely, as in the standard perceptron rule.

Formally, if *ŷ*^*μ*^ = 0 and *y*^*μ*^ = 1 (false negative), we update *w*_*i*_ according to the rule *w*_*i*_ ← *w*_*i*_ + *η*(*w*_*max*_ − *w*_*i*_)*x*_*i*_. If *ŷ*^*μ*^ = 1 and *y*^*μ*^ = 0 (false positive) we update each weight *w*_*i*_ according to the rule *w*_*i*_ + *η*(*w*_*min*_ − *w*_*i*_)*x*_*i*_ (Table 2). Because it is always the case that *w*_*min*_ *< w*_*i*_ *< w*_*max*_, the sign of the weight update in the asymptotic perceptron rule for each case is consistent with the sign of the update in the original perceptron rule; the rules only differ in whether the weights change linearly or asymptotically as a function of the current weight.

**Table 2:**
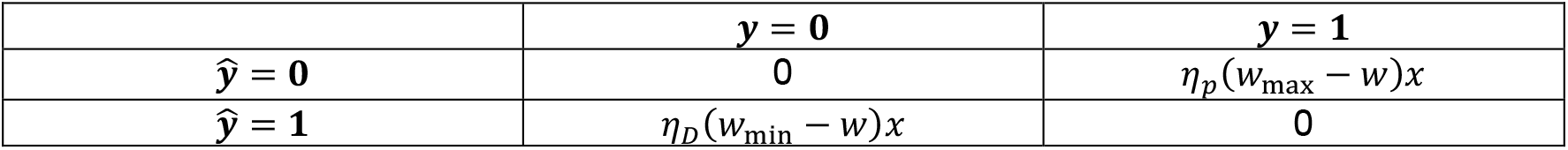
Weight update Δw in the asymptotic perceptron learning rule.

To implement the perceptron learning rule in the calcitron, we again stipulate that the synaptic inputs are binary, so we only have to worry about the direction of synaptic change, not the magnitude. Because the perceptron is a supervised learning algorithm, we will also have a supervisory signal. In our first attempt at implementing the perceptron, the supervisory signal (the “label supervisor”) will simply indicate the value of label (Fig. 8A). In other words, for each input pattern *x*^*μ*^, we have *Z*^*μ*^ = *y*^*μ*^. The challenge here is to set the calcium thresholds and coefficients such that each quadrant of Table 2 (or Table 1 with the standard perceptron rule) is satisfied.

**Figure 8:**
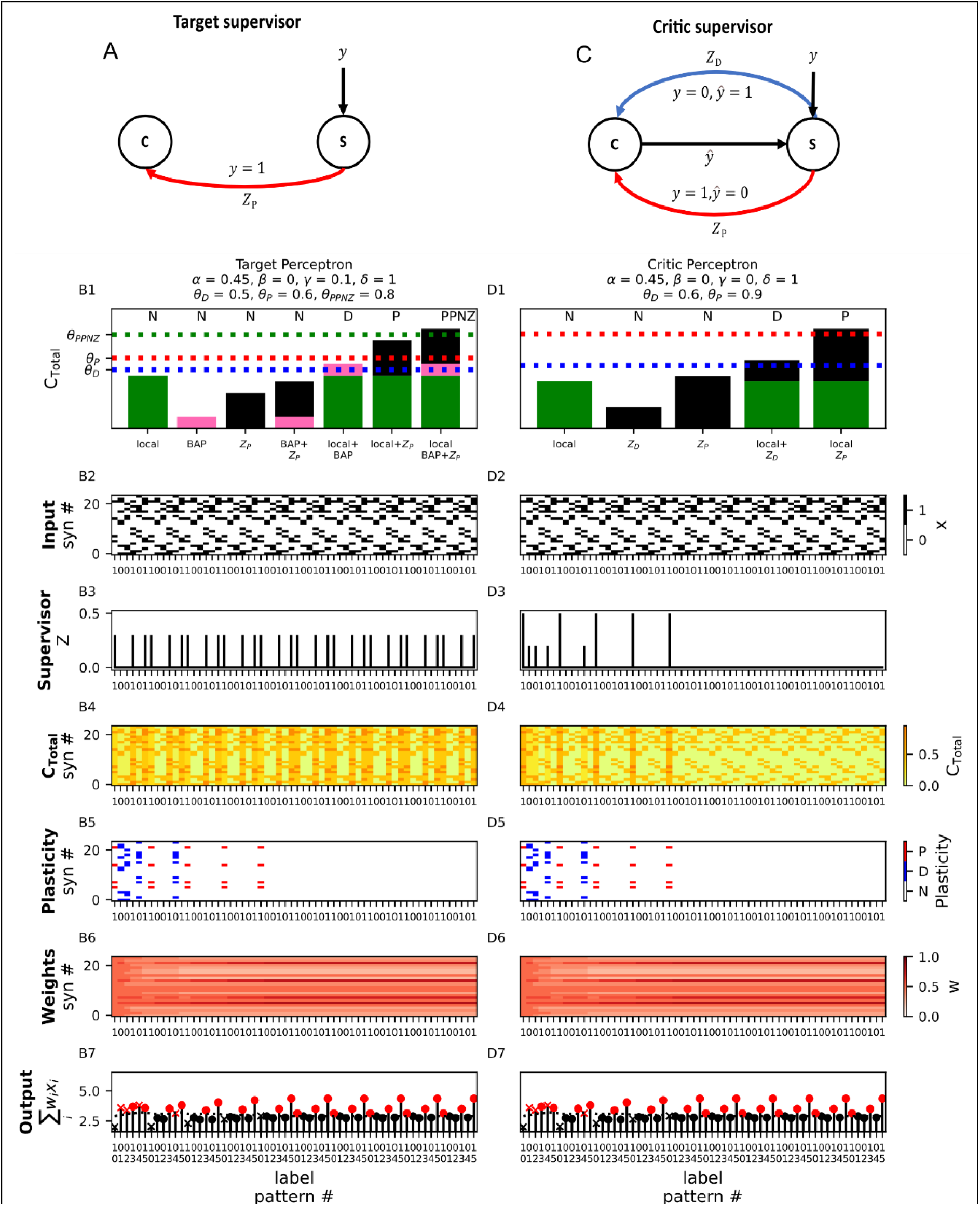
Perceptron learning with the calcitron. **(A)** Supervision circuit for perceptron learning using a “target” supervisor. Whenever the target label is 1, the supervisor sends a potentiative supervisory signal *Z*_P_ to the calcitron. **(B1)** Plasticity rule for the perceptron with a target supervisor. Note that an additional calcium threshold (dashed green line) for a post-potentiative neutral zone (PPNZ) where no plastic change occurs has been added to the plasticity rule. **(B2)** Six patterns, half of which are arbitrarily assigned to the positive class and half to the negative class (*y* = 1 or *y* = 0, respectively, see tick labels on x-axis) are repetitively presented to the calcitron. **(B3)** Supervisory signal. Appears whenever the target label *y* = 1. **(B4-B6)** [Ca^2+^], plasticity, and weights over the course of the simulation. **(B7)** Calcitron output. Red circle: true positive, red ‘x’: false positive, black circle: true negative, black ‘x’: false negative. **(C)** Supervision circuit for perceptron learning using a “critic” supervisor. The supervisor compares the target label *y* to the calcitron output *ŷ*. If the trial was a false negative (*y* = 1, *ŷ*= 0), the supervisor sends a potentiative signal supervisory signal *Z*_P_ to the calcitron. If this trial was a false positive (*y* = 0, *ŷ*= 1), the supervisor sends a depressive supervisory signal *Z*_*D*_ to the calcitron. **(D1-D7)** Perceptron learning with the “critic” supervisory circuit. Note the different magnitudes of the supervisory signal in (D3) – the large signal corresponds to *Z*_P_ and the small signal corresponds to *Z*_*D*_.

To satisfy the upper left quadrant (*ŷ*^*μ*^ = 0 and *y*^*μ*^ = 0), we stipulate that α *<* θ_D_, so that in the absence of a postsynaptic spike and a supervising signal, the presynaptic [Ca^2+^] alone is too low to induce plasticity. For the lower left quadrant, (*ŷ*^*μ*^ = 1 and *y*^*μ*^ = 0, false positives) we require that a postsynaptic spike induces depression at an active synapse, but does nothing to inactive synapses. To accomplish this, we enforce *γ <* θ_D_ and θ_D_ *< γ* + *α <* θ_P_. For the upper right quadrant, (*ŷ*^*μ*^ = 0 and *y*^*μ*^ = 1, false negatives), we want the supervisor in the absence of a postsynaptic spike to induce potentiation at active synapses but do nothing at inactive synapses. For this we require that δ *<* θ_D_ and δ + *α* > θ_P_.

A problem arises, however, when we get to the lower right quadrant (*ŷ*^*μ*^ = 1 and *y*^*μ*^ = 1, true positives). When there is both a postsynaptic spike and a supervising signal, we require that none of the synapses will be updated. But we already enforced that *δ* + *α* > θ_P_, so now when we have both a postsynaptic spike and a supervisory signal we have a [Ca^2+^] of *δ* + *α* + γ at active synapses, which is certainly greater than θ_P_! We solve it by adding a third region of [Ca^2+^], the post-potentiative neutral zone, or PPNZ (θ_PPNZ_ > θ_P_), wherein the [Ca^2+^] is so high that it ceases to potentiate synapses (there is some evidence for this, see (Tigaret et al., 2016)). We then enforce that *δ* + *γ* > θ_PPNZ_, which gives us our fourth quadrant and allows us to reproduce the perceptron learning rule in its entirety (Fig. 8B).

If we don’t wish to add a post-potentiative neutral zone, we can still implement the perceptron learning rule with the normal calcium thresholds if we use a “smart” supervisory signal. Instead of just telling the calcitron what the correct label is, we can have a “critic supervisor” which compares the calcitron’s output *ŷ*^*μ*^ to the target output *y*^*μ*^, and gives supervisory signals of different strengths depending on whether the input pattern resulted in a false positive or false negative. Importantly, the supervisor will no longer give any supervisory signal when the pattern is classified correctly, circumventing the problem we had with true positives using the label supervisor. Here, we will not explicitly describe the circuit necessary to construct such a supervisor, we will simply assume that some other brain circuit can perform the operation of comparing the calcitron’s output to the target output and produce the appropriate supervisory signals (Fig. 8C).

Once we have a critic supervisor, it is no longer necessary to use the calcium from the postsynaptic spike, so we set *γ* = 0. Now the strategy for implementing the perceptron is straightforward. For true positives (*ŷ*^*μ*^ = 1 and *y*^*μ*^ = 1) and true negatives (*ŷ*^*μ*^ = 0 and *y*^*μ*^ = 0), there is no supervisory calcium, so we just have to ensure that an active synapse by itself doesn’t induce plasticity by setting α *<* θ_D_. For false negatives (*ŷ*^*μ*^ = 0 and *y*^*μ*^ = 1) we need the presynaptic calcium combined with the active synapse to induce potentiation, while the supervisor alone doesn’t induce any plasticity at inactive synapses. To do this we set δ = 1 and we construct a potentiation supervisor for false negatives, Z_P_, such that Z_P_ *<* θ_D_ and Z_P_ + α > θ_*P*_. For false positives (*ŷ*^*μ*^ = 1 and *y*^*μ*^ = 0), we similarly construct a supervisor Z_*D*_, such that Z_P_ *<* θ_D_ and θ_*D*_ *<* Z_D_ + α *<* θ_*P*_. These constraints satisfy the rules specified in Table 2 (and Table 1) in a much simpler manner.

To illustrate that the Calcitron can indeed implement the perceptron learning rule just by setting the calcium coefficients and thresholds, we performed a simple classification experiment using a calcitron with the constraints and critic supervisor described above. We generated six binary patterns with 24 synapses each. Half of the patterns were arbitrarily assigned to the positive class and half to the negative class (*y* = 1 or *y* = 0, respectively). We repetitively presented these patterns to both to the “critic” and “target” calcitron supervisory circuits. Initially, the calcitron makes mistakes on some of the patterns, of both the “false positive” and “false negative” variety. After a sufficient number of presentations of each input pattern, however, both supervisory circuits succeed to ensure that the calcitron correctly classifies all patterns (Fig. 8).

We have thus demonstrated that the perceptron learning algorithm can be implemented with the calcitron, either with a “label supervisor” – which requires an additional plasticity threshold, or with a “critic supervisor” which can use the standard plasticity thresholds.

## Discussion

### Summary

We have shown that the calcitron, a simple model neuron that uses four different sources of calcium to modify its synaptic weights, can implement a wide variety of learning and plasticity rules. We have demonstrated that merely by appropriately setting the amount of calcium influx synapses receive from each calcium source, it is possible to reproduce classic learning rules, like Hebb’s rule (Fig. 2) or the perceptron learning rule (Fig. 8). We have also reproduced, in a simplified manner, plastic phenomena observed in biology, such as frequency-dependent plasticity (Fig. 4, (O’Connor et al., 2005b)), homeostatic plasticity (Fig. 7, (G. Turrigiano, 2012; G. G. Turrigiano, 2008; G. G. Turrigiano & Nelson, 2004)), and behavioral time-scale plasticity (Bittner et al., 2015, 2017; Grienberger & Magee, 2022; Milstein et al., 2021). Moreover, we have shown how the calcium control hypothesis can result in novel plasticity and learning rules, such as the 14 pre-post rules (Fig. 3) and the unsupervised learning of large repetitive patterns with heterosynaptic plasticity (Fig. 5). We note that because each calcium source in the Calcitron is related to a current source (e.g. the presynaptic input or postsynaptic spike), the Calcitron bears some similarity to the BCM plasticity rule (Bienenstock et al., 1982).

The calcium control hypothesis was originally developed as an explanation for Hebbian and anti-Hebbian plasticity (J. Lisman, 1989). And early mathematical formulations of the calcium control hypothesis were designed to reproduce frequency-dependent and spike timing-dependent plasticity (Graupner & Brunel, 2012; Shouval et al., 2002). With the calcitron, however, we have expanded these earlier results into a generalized, simple model which can explain and predict a much wider set of plasticity and learning results from first principles. The calcitron can thus be helpful in providing an intuition for the range of possible calcium-based mechanisms underlying experimentally-observed plasticity, including forms of plasticity that are as-yet undiscovered.

The mathematical formalism of the calcitron also makes it easier to understand the limitations of calcium-based plasticity. The calcitron equations impose constraints on the potential learning rules that can emerge from the calcium control hypothesis, which can help us determine whether it is necessary to posit additional mechanisms, such as the supervisory circuit we proposed for homeostatic plasticity (Fig. 7). Importantly, we do not claim that every possible learning rule we propose here is implemented in biology as we have described it. Rather, the calcitron is intended to serve as a framework for exploring different predictions of the calcium control hypothesis.

### Future Directions and Additional Biological Considerations

The calcitron model was formulated to be as simple as possible in order to provide straightforward mathematical intuitions about the calcium basis of synaptic plasticity. This simplicity comes with certain drawbacks. For example, the choice of using a perceptron-like neuron model that does not temporally integrate information means that the calcitron, as formulated here, cannot implement spike timing-dependent plasticity (STDP). Prior work has shown that a leaky integrator model with calcium dynamics can indeed generate STDP (Graupner & Brunel, 2012; Shouval et al., 2002). As such, future work can consider leaky integrator versions of the calcitron that can implement STDP and other temporally-sensitive plasticity rules. However, this temporal sensitivity introduces the need to finely tune calcium decay time constants in order to produce different plasticity rules, which introduces a level of complexity beyond the scope of the current work. It is also possible to explicitly model the kinetics of the relevant ion channels (i.e. VGCCs and NMDA receptors), adding even more biological detail at the expense of greater complexity.

The calcitron is also linear, both in terms of its input-output function (excluding the activation function) and in terms of how calcium from different sources combines to make calcium. Linear point neurons are commonly used to model neural phenomena, although experimental and theoretical work indicate that real neurons may integrate information in a nonlinear fashion (Beniaguev et al., 2021; Gordon et al., 2006; Mel, 1991; Moldwin et al., 2021; Moldwin & Segev, 2020; Poirazi et al., 2003; Poirazi & Mel, 2001; Polsky et al., 2004, 2009; Schiller et al., 2000; Tran-Van-Minh et al., 2015). Of particular relevance is the superlinear activation function of the NMDA receptor, whose conductance exhibits sigmoidal sensitivity to local voltage at the synapse location (Jahr et al., 1990; Jahr & Stevens, 1990). A more realistic model for local calcium influx could thus include this voltage-dependent nonlinearity, in line with experimental work showing that NMDA spikes induce plasticity (Kumar et al., 2021).

Another important feature of biological neurons not incorporated into the calcitron is the spatial distribution of synapses on a neuron’s dendrites. This is especially relevant for heterosynaptic plasticity, which depends on the location of synapses relative to each other as well as their absolute location on the dendritic tree (Chater & Goda, 2021; Chistiakova et al., 2014; Moldwin et al., 2022; Tong et al., 2021). The calcitron can be augmented to include location-dependent heterosynaptic plasticity on a single dendrite following the schemes of the clusteron (Mel, 1991) or the G-clusteron (Moldwin et al., 2021) or to explicitly include a branching dendrite to account for the branch-dependent hierarchical heterosynaptic plasticity effect we posited in previous work (Moldwin et al., 2022). The spatial structure of the dendrite can also influence homeostatic plasticity (Rabinowitch & Segev, 2006a, 2006b) as well as how inhibitory inputs affect plasticity at excitatory synapses (Bar-Ilan et al., 2012).

A crucial assumption we made in formulating the calcitron is that local calcium is always a function of presynaptic activity. If we had allowed for synapse-specific supervisory signals, it would be possible to implement arbitrary learning rules without any concern for constraints, as we could simply engineer a supervisor to potentiate or depress each synapse independently. In biology, however, it is possible that synapse-specific plasticity supervisors exist. One candidate for such a mechanism are the internal calcium stores in the endoplasmic reticulum (ER) (Rose & Konnerth, 2001). The ER can invade individual spines (Spacek & Harris, 1997), making it a candidate for targeted calcium release at selected synapses. It is usually assumed that calcium released from the ER is mediated by calcium influx via a process called calcium-induced calcium release, or CICR. If the initial calcium needed to induce CICR at the ER in individual spines come from extracellular sources, CICR is still broadly consistent with our model; the internal calcium sources may simply be amplifying the calcium which enters via external sources. If, however, there are other endogenous mechanisms that can differentially induce calcium release from the ER at different synapses, the ER may be able to play the role of a synapse-specific supervisor. An additional candidate for synapse-specific supervisors are astrocytes, which target individual dendritic spines (Haber et al., 2006) and can influence plasticity via a variety of signaling pathways (Barker & Ullian, 2010). Either of these mechanisms can enable a less constrained, more diverse palette of neural learning rules when combined with the more classical calcium sources we suggested here.

With respect to the supervisory signal, we have focused on plasticity supervisors that operate via calcium mechanisms, such as observed in Purkinje neurons (Konnerth et al., 1992) and possibly in the CA1 region of the hippocampus during BTSP induction. However, there are other forms of plasticity that are more dependent on neuromodulation. For example, associative learning is often thought to require dopamine as a supervisory signal (Lisman et al., 2011; Puig et al., 2014). Future work can explore the interplay between calcium-based and neuromodulator-dependent mechanisms of plasticity.

There are a variety of ways to build on the calcitron. In this work, we have focused on building neurons and circuits that implement one learning rule at a time. It may be desirable, however, for a neuron to simultaneously implement, for example, Hebbian plasticity and homeostatic plasticity. The calcitron equations constrain which kind of learning rules can be simultaneously implemented in this fashion. It would be worthwhile to explore which learning rule combinations are mathematically valid in the calcitron framework.

Another natural extension of the calcitron would be to incorporate multiple calcitron nodes into a network to see how calcium-based plasticity affect network activity. It would be interesting to see how networks containing neurons with different calcium coefficients and thresholds (and thus different learning rules) would operate. Alternatively, it is possible to build large networks containing subnetworks with different calcium parameters, to explore how brain regions with different plasticity rules might interact with each other. The calcitron framework thus provides a theoretical tool to explore the network-level consequences of biologically realistic plasticity rules.

## Methods

All simulations were performed on a Windows PC using Python and plotted using the Matplotlib library (Hunter, 2007). Parameters for each simulation are given in the table below:

| Fig | Pane<br>l | $\eta_D$ | $F_D$ | $\theta_P$ | $F_P$ | $\eta_P$ | $\alpha$ | $\beta$ | $\gamma$ | $\delta$ | $b$ | $g$ | $N$ | $Z$ | $Z_P$ | $Z_D$ | $\eta_{PPNZ}$ | $F_{PPNZ}$ | $\eta_b$ |
| --- | --- | --- | --- | --- | --- | --- | --- | --- | --- | --- | --- | --- | --- | --- | --- | --- | --- | --- | --- |
| Fig<br>1 |  |  |  |  |  |  |  |  |  |  |  |  |  |  |  |  |  |  |  |
|  | E-G | -0.1 |  | 1 |  | 0.2 |  |  |  |  |  |  |  |  |  |  |  |  |  |
|  | H-L | 0.1 | 0 | 1 | 1 | 0.1 |  |  |  |  |  |  |  |  |  |  |  |  |  |
| Fig<br>2 |  |  |  |  |  |  |  |  |  |  |  |  |  |  |  |  |  |  |  |
|  | A1-F1 | 0.1 | 0 | 0.8 | 1 | 0.1 | 0.4 | 0 | 0.4<br>5 | 0 | -<br>1.<br>8 | step | 10 |  |  |  |  |  |  |
|  | A2-F2 | 0.1 | 0 | 0.8 | 1 | 0.1 | 0.5<br>5 | 0 | 0.7 | 0 | -<br>1.<br>8 | step | 10 |  |  |  |  |  |  |
|  | A3-F3 | 0.1 | 0 | 0.8 | 1 | 0.1 | 0.4 | 0 | 0.3 | 0 | -<br>1.<br>8 | step | 10 |  |  |  |  |  |  |
|  | A4-F4 | 0.1 | 0 | 0.5 | 1 | 0.1 | 0.5<br>5 | 0 | 0.6 | 0 | -<br>1.<br>8 | step | 10 |  |  |  |  |  |  |
| Fig<br>3 |  |  |  |  |  |  |  |  |  |  |  |  |  |  |  |  |  |  |  |
|  | 1,1 | 1 | 0 | 0.8 | 1 | 1 | 0.2 | 0 | 0.2 | 0 |  |  |  |  |  |  |  |  |  |
|  | 1,2 | 1 | 0 | 0.8 | 1 | 1 | 0.3 | 0 | 0.3 | 0 |  |  |  |  |  |  |  |  |  |
|  | 1,3 | 1 | 0 | 0.8 | 1 | 1 | 0.1 | 0 | 0.6 | 0 |  |  |  |  |  |  |  |  |  |
|  | 1,4 | 1 | 0 | 0.8 | 1 | 1 | 0.6 | 0 | 0.1 | 0 |  |  |  |  |  |  |  |  |  |
|  | 2,1 | 1 | 0 | 1.3 | 1 | 1 | 0.6 | 0 | 0.6 | 0 |  |  |  |  |  |  |  |  |  |
|  | 2,2 | 1 | 0 | 0.8 | 1 | 1 | 0.4<br>5 | 0 | 0.4<br>5 | 0 |  |  |  |  |  |  |  |  |  |
|  | 2,3 | 1 | 0 | 0.8 | 1 | 1 | 0.3 | 0 | 0.6 | 0 |  |  |  |  |  |  |  |  |  |
|  | 2,4 | 1 | 0 | 0.8 | 1 | 1 | 0.3 | 0 | 0.9 | 0 |  |  |  |  |  |  |  |  |  |
|  | 3,1 | 1 | 0 | 0.8 | 1 | 1 | 0.6 | 0 | 0.3 | 0 |  |  |  |  |  |  |  |  |  |
|  | 3,2 | 1 | 0 | 0.8 | 1 | 1 | 0.6 | 0 | 0.6 | 0 |  |  |  |  |  |  |  |  |  |
|  | 3,3 | 1 | 0 | 0.8 | 1 | 1 | 0.6 | 0 | 0.9 | 0 |  |  |  |  |  |  |  |  |  |
|  | 3,4 | 1 | 0 | 0.8 | 1 | 1 | 0.9 | 0 | 0.3 | 0 |  |  |  |  |  |  |  |  |  |
|  | 4,1 | 1 | 0 | 0.8 | 1 | 1 | 0.9 | 0 | 0.6 | 0 |  |  |  |  |  |  |  |  |  |
|  | 4,2 | 1 | 0 | 0.8 | 1 | 1 | 0.9 | 0 | 0.9 | 0 |  |  |  |  |  |  |  |  |  |
| Fig<br>4 |  |  |  |  |  |  |  |  |  |  |  |  |  |  |  |  |  |  |  |
|  | A | 0.1 | 0 | 0.8 | 1 | 0.1 | 0.3 | 0 | 0.3 | 0 |  |  |  |  |  |  |  |  |  |
|  | B1 | 0.1 | 0 | 0.8 | 1 | 0.1 | 0.3 | 0 | 0 | 0 |  |  |  |  |  |  |  |  |  |
|  | B2 | 0.1 | 0 | 0.8 | 1 | 0.1 | 0.3 | 0 | 0.1<br>5 | 0 |  |  |  |  |  |  |  |  |  |
|  | B3 | 0.1 | 0 | 0.8 | 1 | 0.1 | 0.3 | 0 | 0.3 | 0 |  |  |  |  |  |  |  |  |  |
|  | B4 | 0.1 | 0 | 0.8 | 1 | 0.1 | 0.1<br>5 | 0 | 0.3 | 0 |  |  |  |  |  |  |  |  |  |
|  | B5 | 0.1 | 0 | 0.8 | 1 | 0.1 | 0 | 0 | 0.3 | 0 |  |  |  |  |  |  |  |  |  |
| Fig<br>5 |  |  |  |  |  |  |  |  |  |  |  |  |  |  |  |  |  |  |  |
|  | All | 0.02<br>5 | 1 | 2.6 | 3 | 0.02<br>5 | 2 | 0.<br>2 | 0 | 0 | -8 | step | 12 |  |  |  |  |  |  |
| Fig<br>6 |  |  |  |  |  |  |  |  |  |  |  |  |  |  |  |  |  |  |  |
|  | All | 1 | 0 | 0.7 | 1 | 1 | 0.2 | 0 | 0 | 0.<br>6 | -6 | step | 14 | 1 |  |  |  |  |  |
| Fig<br>7 |  |  |  |  |  |  |  |  |  |  |  |  |  |  |  |  |  |  |  |
|  | B | 0.02 | 0 | 1 | 0.<br>5 | 0.02 | 0 | 0 | 0.2 | 1 | 0 | linea<br>r | 20 |  | 1.0<br>5 |  |  |  |  |
|  | C | 0.02 | 0 | 0.7 | 0.<br>5 | 0.02 | 0.3<br>5 | 0 | 0.0<br>6 | 1 | 0 | linea<br>r | 20 |  | 0.3<br>5 |  |  |  |  |
|  | E | 0.02 | 0 | 1 | 0.<br>5 | 0.02 | 0 | 0 | 0 | 1 | 0 | linea<br>r | 20 |  | 1.0<br>5 | 0.5<br>5 |  |  |  |
|  | F | 0.02 | 0 | 0.7 | 0.<br>5 | 0.02 | 0.3 | 0 | 0 | 1 | 0 | linea<br>r | 20 |  | 0.4<br>5 | 0.2<br>5 |  |  |  |
| Fig<br>8 |  |  |  |  |  |  |  |  |  |  |  |  |  |  |  |  |  |  |  |
|  | B | 0.2 | 0 | 0.6 | 1 | 0.2 | 0.4<br>5 | 0 | 0.1 | 1 | -<br>2.<br>8 | step | 24 |  | 0.3 |  | 0 | 0.25 | 0.1 |
|  | D | 0.2 | 0 | 0.9 | 1 | 0.2 | 0.4<br>5 | 0 | 0 | 1 | -<br>2.<br>8 | step | 24 |  | 0.5 | 0.2 |  |  | 0.1 |

## Code Availability

Code for this project can be found at https://github.com/tmoldwin/Calcitron.

